# Reducing Glucocorticoid Burden in Lupus with Omega-3 Fatty Acids: Docosahexaenoic Acid Augments Prednisone Efficacy in Maintaining Cyclophosphamide-Induced Remission of Preclinical Lupus Nephritis

**DOI:** 10.64898/2026.01.07.698195

**Authors:** Ashley N. Anderson, Olivia F. McDonald, Shayla-Rae S. Johnson, John J. Liddle, Vanessa Estrada, Jalen T. Jackson, Ryan P. Lewandowski, James G. Wagner, Jack R. Harkema, James J. Pestka

**Author notes:** **Corresponding Author:** Dr. J.J. Pestka, Department of Microbiology, Genetics, and Immunology, Michigan State University, East Lansing, MI, United States.

## Abstract

**Background:** Managing lupus nephritis (LN) remains challenging due to relapses after immunosuppressive induction and toxicity from long-term glucocorticoid (GC) maintenance therapy. Dietary omega-3 fatty acids prevent LN onset in preclinical models, but their role in maintaining remission post-induction remains unstudied.

**Methods:** The silica-accelerated LN (SALN) model using lupus-prone NZBWF1 mice was used to evaluate how docosahexaenoic acid (DHA), an omega-3 fatty acid, influenced LN remission durability after cyclophosphamide (CYC) induction, alone or with a moderate dose of prednisone (PDN). Mice received intranasal silica weekly from 8 to 11 weeks. After LN developed at 21 weeks, groups were injected weekly with CYC (human equivalent dose [HED]=31 mg/day) or vehicle (VEH) for 8 weeks, during which CYC groups also received control, DHA (HED=5 g/day), PDN (HED=9 mg/day), or DHA+PDN diets. Disease activity was monitored via proteinuria, autoantibodies, and survival. Six weeks post-CYC, multi-organ histopathology and immunohistochemistry were assessed.

**Results:** VEH-treated mice developed severe LN with early death. CYC slowed disease temporarily in control- and PDN-fed mice; relapses occurred after cessation. DHA or DHA+PDN increased tissue omega-3 levels and prolonged remission. PDN and DHA monotherapies and co-therapy improved survival, but DHA+PDN was most effective at sustaining remission as reflected by reduced histopathologic markers of lupus severity in the kidney, spleen, lung, and brain.

**Conclusion:** DHA+PDN optimally maintained LN remission after CYC, supporting omega-3 supplementation as a potential GC-sparing strategy to improve immunosuppressive therapy and prevent relapses.

**Highlights:**

- Frequent post-immunosuppressive treatment flares and the toxicity from long-term maintenance therapy with glucocorticoids (GCs) limit effective management of lupus nephritis (LN).
- The silica-accelerated lupus nephritis (SALN) model in NZBWF1 mice mimics the gene–environment interactions of human systemic lupus erythematosus (SLE) and LN, enabling synchronized and efficient preclinical studies of drug and nutritional interventions.
- Short-term immunosuppressive therapy with cyclophosphamide (CYC) induces temporary remission of LN and extrarenal inflammation and autoimmunity in SALN mice, which can be extended through monotherapy with either dietary supplementation with omega-3 docosahexaenoic acid (DHA) or a moderate dose of the GC prednisone (PDN).
- Combined maintenance therapy with DHA and PDN proved more effective than monotherapy in enhancing the durability of post-CYC LN remission and in reducing extrarenal inflammation and autoimmunity in the lung, spleen, and brain.
- These preclinical findings show that dietary supplementation with omega-3s such as DHA may offer a safe, affordable, GC-sparing adjunctive option for managing LN and SLE.

## Introduction

Systemic lupus erythematosus (SLE) is a severe, multifactorial autoimmune disease that causes immune-mediated damage across multiple organs, including the skin, spleen, lungs, brain, and kidneys [1–4]. Its onset and progression are driven by complex gene–environment interactions and characterized by recurrent flares that substantially increase morbidity and the risk of irreversible organ damage. Lupus nephritis (LN) is a major complication that can progress to end-stage renal disease when inadequately controlled [1,4,5].

Conventional therapy for advanced proliferative LN (Class III or higher) typically follows a two-phase paradigm to quell renal inflammation, suppress autoimmunity, and promote durable remission [6]. The induction phase uses potent immunosuppressants such as cyclophosphamide (CYC) to achieve LN remission, followed by a maintenance phase to sustain remission and limit cumulative damage, using tapered doses of glucocorticoids (GCs) like prednisone (PDN) together with other drugs, such as mycophenolate mofetil [7,8]. Although these regimens are effective, GCs have myriad adverse effects—including serious infections, fatigue, metabolic and cardiovascular complications, type 2 diabetes, bone loss, reproductive and hepatic toxicity, and bone marrow suppression—that pose ongoing challenges in selecting drugs, determining treatment duration, and designing tapering strategies. Importantly, prematurely reducing or stopping maintenance therapy significantly increases the risk of relapse and flaring [9]. Recent advances in biologic and targeted therapies have expanded the LN armamentarium and can theoretically reduce GC exposure [6], yet their high cost and restricted availability remain significant barriers for the ever-growing population of patients without adequate insurance coverage [10].

There is increasing recognition that dietary modulation has value as a complementary strategy for treating persistent inflammatory and autoimmune diseases, an emphasis reflected in initiatives such as the NIH Strategic Blueprint on Precision Nutrition and the NIH-Wide Strategic Plan for Autoimmune Disease Research [11,12]. Consistent with this recognition, many clinicians are adopting dietary counseling into routine SLE care [13], and there is particular interest in precision nutrition approaches that target the tissue lipidome by increasing consumption of omega-3 polyunsaturated fatty acids (PUFAs) [14–16]. Western dietary patterns, typically enriched in omega-6 PUFAs such as linoleic acid (LA) from seed oils and arachidonic acid (ARA), formed by LA elongation and found in animal fat, are associated with worsening of chronic inflammatory states driven by ARA-derived metabolites [17]. In contrast, diets or supplements rich in marine-derived omega-3 PUFAs—especially docosahexaenoic acid (DHA) and eicosapentaenoic acid (EPA), sourced from microalgae, krill, or cold-water fish oils—can attenuate inflammatory responses and promote their resolution [18].

A well-established literature links DHA and EPA intake of 1 to 5 g/day to favorable changes in cardiovascular risk biomarkers and outcomes, as well as clinical improvement in inflammatory rheumatologic conditions [19–23]. Regulatory investigations confirm that combined EPA+DHA supplementation up to 5 g/day is safe for adults [24]. This dosage costs approximately $2/day when purchased over the counter as a stable, high-quality supplement [25]. Mechanistically, omega-3s are rapidly incorporated into immune cell membranes and act by i) altering membrane fluidity and lipid raft organization to modulate receptor clustering and downstream signaling, ii) dampening activation of proinflammatory transcription factors such as NF-κB while activating others with anti-inflammatory actions like PPAR-γ, iii) reducing proinflammatory gene expression and cytokine production, iv) suppressing production of inflammatory ARA-derived leukotrienes, prostaglandins, and thromboxanes by competitive inhibition, and v) serving as substrates for specialized pro-resolving mediators that drive active resolution of inflammation and restoration of tissue homeostasis [15,18,26]. Remarkably, many of these molecular targets overlap with or complement those of GCs [27], offering promise of safe adjunctive omega-3 supplementation to reduce flare frequency and slow kidney damage in individuals with lupus.

Preclinical investigations in lupus-prone mouse strains over four decades have consistently shown that omega-3-enriched diets suppress or delay autoimmune responses, reduce the onset and progression of LN, and improve survival rates [28–53]. In contrast, diets high in saturated fats, monounsaturated fats, and omega-6s, typical of ultraprocessed foods, have been linked to increased autoimmunity and kidney damage. Building upon this base, our lab has developed an efficient gene–environment interaction model in autoimmune-prone NZBWF1 mice, in which short-term, repeated intranasal instillations of crystalline silica (cSiO_2_), a well-known environmental trigger of lupus, accelerate systemic autoimmunity and LN by approximately three months, providing a highly reproducible silica-accelerated LN (SALN) platform [54,55]. Within this model, oral supplementation with microalgal oil–derived DHA at human equivalent doses (HEDs) of 2 to 5 g/day provides broad dose-dependent protection, including: i) suppression of inflammatory, type I interferon, and autoimmune gene networks in the lung, ii) reduction of B- and T-cell infiltration and ectopic lymphoid structure (ELS) formation, iii) lowering of pulmonary and systemic autoantibody (AAb) titers, iv) prevention of glomerulonephritis, and v) prolonging survival [26,56–63]. Collectively, these studies position DHA supplementation as a suitable adjunctive strategy for reducing GC use during LN treatment.

The SALN model has been further leveraged to assess the preventive effectiveness of the PDN, demonstrating that oral pretreatment with a medium PDN dose (14 mg/day HED) substantially reduced cSiO_2_-induced pulmonary ELS formation, AAb production, and inflammatory and autoimmune gene expression in the lung and kidney. This dose also reduced splenomegaly and the severity of glomerulonephritis [64]. In contrast, low-dose PDN (5 mg/day HED) did not improve autoimmune symptoms, whereas a high-dose PDN (46 mg/day HED) caused severe GC-associated toxicity, leading to early cohort termination. Significantly, even at the effective medium dose, PDN caused muscle wasting and did not improve survival. This indicates a discordance between histopathologic improvements and death rates in SALN and confirms the narrow therapeutic window of long-term GC therapy [64]. Furthermore, these observations suggest that, within the constraints of the SALN model, DHA offers a more favorable balance between preventive effectiveness and tolerability than PDN.

Although a substantial amount of preclinical data supports the use of omega-3s in lupus, including research with DHA in SALN, these studies have primarily focused on prevention rather than treatment of established disease. This limits their relevance for patients with SLE who often present with active LN that requires remission induction. To address this gap, the SALN model was used here to assess whether adjunctive DHA supplementation can maintain LN remission after immunosuppressive induction with CYC, when administered either alone or combined with a moderate, clinically relevant dose of PDN. We show for the first time preclinical evidence that adjunctive DHA improves the efficacy of PDN, prolonging remission of both LN and extrarenal lupus pathogenesis. Accordingly, safe, inexpensive omega-3 supplementation might serve as a GC-sparing strategy to improve long-term disease management and reduce treatment-related toxicity in LN and SLE.

## Materials and Methods

### Animals

All experimental protocols were reviewed and approved by the Institutional Animal Care and Use Committee (IACUC) at Michigan State University in accordance with the NIH guidelines (AUF #PROTO202400258). Female 7-week-old lupus-prone NZBWF1 mice were obtained from the Jackson Laboratory (#100008, Bar Harbor, ME), randomly assigned four per cage with free access to food and water, and kept at a consistent temperature (20–24°C) and humidity (30–70%) with a 12-hour light/dark cycle.

### Diets

Diet formulations were based on the nutritionally balanced semi-purified American Institute of Nutrition (AIN)-93G rodent diet containing 70 g soybean oil per kg of diet (AIN-93G rodent diet #110700, Dyets Inc., Bethlehem, PA) [65]. Four experimental diets were used in the study (Table 1). The (i) basal control diet (CON) was made by replacing the 70 g/kg soybean oil with 60 g/kg of high-oleic safflower oil (AVO high-oleic safflower oil, American Vegetable Oils, Inc., Commerce, CA) and 10 g/kg of corn oil (Mazola Corn Oil, ACH Food Companies, Inc., Oakbrook Terrace, IL) to supply essential fatty acids. For the (ii) DHA diet, 25 g/kg of high-oleic safflower oil was substituted in the basal CON diet with 25 g/kg of microalgal oil containing 40% DHA (DHASCO, DSM Nutritional Products, Columbia, MD), as described earlier [56,57]. This diet contained 10 g/kg DHA, modeling human DHA intake of 5 g per day on a caloric basis (Table 1). For a patient, this would be roughly equivalent to taking i) five triple-strength over-the-counter fish oil capsules daily, each containing approximately 1 g of omega-3 fatty acids or ii) five prescription-grade omega-3 capsules used to treat hypertriglyceridemia. For the (iii) PDN diet and the (iv) combined DHA+PDN diet, PDN was dissolved in a small amount of 100% (v/v) ethanol and added to both the basal CON and DHA diets at 10 mg/kg, as previously described [64]. This dietary level corresponds to an HED of 9.4 mg/day (Table 2) and was designated as “moderate” because it falls within the lower end of the medium dose range (7.5 to 30 mg/d) used in GC treatment of rheumatic diseases [66].

**Table 1.** Formulation of experimental diets.

| Macronutrients<sup>a</sup> | Experimental Diets |  |  |  |
| --- | --- | --- | --- | --- |
|  | g/kg Total Diet |  |  |  |
|  | CON | DHA | PDN | DHA+PDN |
| **Carbohydrates** |  |  |  |  |
| Corn Starch | 397.486 | 397.486 | 397.486 | 397.486 |
| Maltodextrin (Dyetrose) | 132 | 132 | 132 | 132 |
| Sucrose | 100 | 100 | 100 | 100 |
| Cellulose | 50 | 50 | 50 | 50 |
| **Proteins** |  |  |  |  |
| Casein | 200 | 200 | 200 | 200 |
| L-Cysteine | 3 | 3 | 3 | 3 |
| **Fats** |  |  |  |  |
| Corn Oil<sup>b</sup> | 10 | 10 | 10 | 10 |
| High-Oleic Safflower Oil<sup>c</sup> | 60 | 35 | 60 | 35 |
| DHA-enriched Algal Oil<sup>d</sup> | 0 | 25 | 0 | 25 |
| **Essential nutrients** |  |  |  |  |
| AIN-93G Mineral Mix | 35 | 35 | 35 | 35 |
| AIN-93G Vitamin Mix | 10 | 10 | 10 | 10 |
| Choline Bitartrate | 2.5 | 2.5 | 2.5 | 2.5 |
| **Preservatives** |  |  |  |  |
| TBHQ Antioxidant | 0.014 | 0.014 | 0.014 | 0.014 |
| **Drug** |  |  |  |  |
| Prednisone | 0 | 0 | 0.01 | 0.01 |

Values in table are shown as mass in g/kg diet. Abbreviations: CON: control, DHA: docosahexaenoic acid, PDN: prednisone. <sup>a</sup>Diet composition is based on the manufacturer's description [65]. <sup>b</sup>Corn oil contributes 6 g/kg diet of linoleic acid and 3 g/kg diet of oleic acid. <sup>c</sup>High-oleic safflower oil contributes 47 g/kg diet of oleic acid and 9 g/kg diet of linoleic acid (https://fdc.nal.usda.gov/). <sup>d</sup>Algal oil contributes 10 g/kg diet of DHA and 4 g/kg diet of oleic acid [114]. A 10 g/kg diet of DHA is calorically equivalent to a human equivalent dose of 5 g/d.

**Table 2.** Relation of mouse cyclophosphamide (CYC) injections and dietary prednisone (PDN) to human doses.

| **Drug** | **Diet concentration**<br>**(mg/kg)** | **Mouse dose**<br>**(mg/kg bw/d)** | **HED**<br>**(mg/kg bw/d)<sup>b</sup>** | **HED**<br>**(mg/d)<sup>c</sup>** |
| --- | --- | --- | --- | --- |
| CYC | 0 | 5 | 0.4 | 31.2 |
| PDN | 10 | 1.5<sup>a</sup> | 0.12 | 9.36 |

<sup>a</sup>Mouse dose was calculated by multiplying dietary concentration by the FDA conversion factor of 0.15 [115].
<sup>b</sup>Human equivalent dose (HED) was calculated by multiplying mouse dose by FDA drug interspecies conversion factor of 0.08 [116].
<sup>c</sup>Daily human equivalent dose based on average body weight of 78 kg for women $\geq 20$ years of age in the U.S. [117].

### Crystalline silica (cSiO_2_)

cSiO_2_ (Min-U-Sil-5, 1.5–2.0 μm average particle size, U.S. Silica, Katy, TX) was acid-washed in 1 M HCl at 100°C for 1 hour to remove surface contaminants that could obscure surface activity. The cSiO_2_ was then rinsed three times with sterile water and dried in an oven at 200°C to eliminate water, endotoxins, and microbial contaminants. On each day of cSiO_2_ instillation, a working stock suspension of 40 mg/ml in Dulbecco’s phosphate-buffered saline (DPBS) was prepared, then sonicated for 1 minute using the Q125 Qsonica sonicator (Qsonica LLC, Newtown, CT) set to 2 minutes with pulses of 5 seconds on and 5 seconds off at 35% amplitude. After sonication, the suspension was inverted and vortexed continuously during instillations to prevent cSiO_2_ settling.

### Experimental design

The experimental study design is depicted in Figure 1. NZBWF1 mice (7 weeks old) were started on the CON diet, and one week later, they were anesthetized with 4% isoflurane and intranasally instilled with 1.0 mg of cSiO_2_ in 25 μl of DPBS once a week for 4 weeks [54]. The total cSiO_2_ dose (4 mg) approximately equals half of a human lifetime exposure according to guidelines set by the Occupational Safety and Health Administration (OSHA)[67]. At 20 weeks of age, cSiO_2_-treated mice were divided into weight-based quartiles, with 1 mouse from each quartile housed together, totaling 4 mice per cage. This grouping ensured balanced experimental conditions, reducing confounding effects from body weight differences. At 21 weeks, when cSiO_2_-treated mice exhibit proteinuria, cages were split into five treatment groups: (i) weekly DPBS vehicle (VEH) intraperitoneal (IP) injections with the CON diet (cSiO_2_/VEH/CON), (ii) weekly CYC IP injections at 35 mg/kg body weight (bw) with the CON diet (cSiO_2_/CYC/CON), (iii) weekly CYC IP injections with a DHA diet (cSiO_2_/CYC/DHA), (iv) weekly CYC IP injections with a PDN diet (cSiO_2_/CYC/PDN), and (v) weekly CYC IP injections with a DHA+PDN diet (cSiO_2_/CYC/DHA+PDN) (Figure 1). To reduce lipid oxidation products, diets were prepared every two weeks, stored at −20°C until used, and provided fresh to the mice daily. Injections with VEH or CYC ceased after 8 weeks, but mice remained on their respective maintenance diets for the remainder of the study. Urine was collected weekly and tested for protein using reagent dipsticks (Siemens Uristix 4, Malvern, PA). Plasma samples were collected biweekly, starting at 27 weeks of age, for blood urea nitrogen (BUN) and AAb analyses. Mice were monitored daily for clinical signs of moribundity (Supplementary Table 2) during feedings, and their body weights were recorded weekly. Any mice meeting moribund criteria were immediately euthanized and necropsied. The study concluded at age 35 weeks with a full necropsy of all remaining animals.

**Figure 1.**
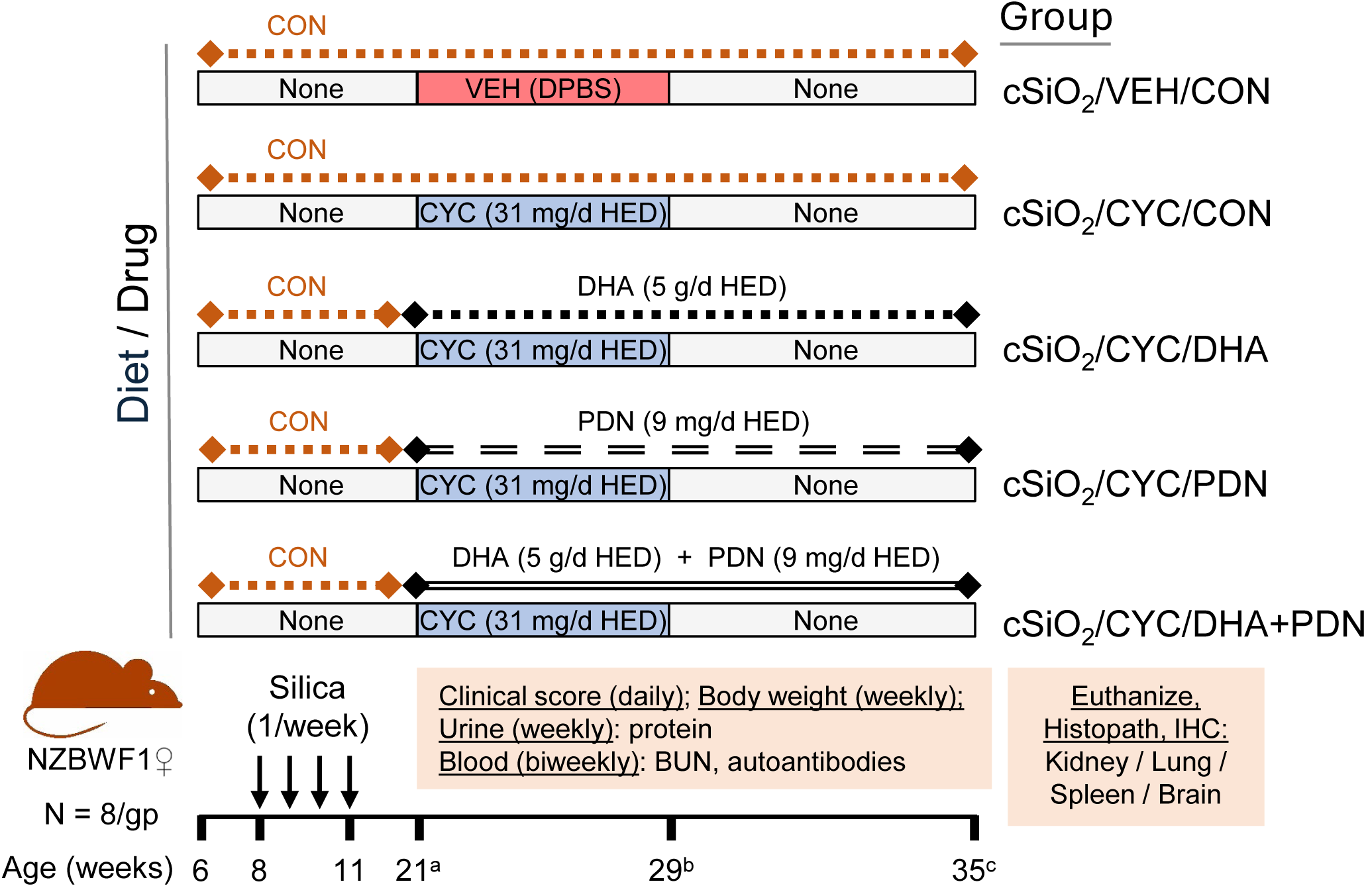
Experimental design for determining how diets containing docosahexaenoic acid (DHA) and/or prednisone (PDN) impact the duration of cyclophosphamide (CYC)-induced remission in the silica (cSiO_2_)-accelerated lupus nephritis (SALN) mouse model. Abbreviations: CON, control diet; VEH, vehicle. Letters: a) nephritis onset/initiate vehicle (VEH) or CYC injections; b) CYC discontinued; c) experiment terminated and mice necropsied when ≥50% CYC-treated CON-fed mice exhibited proteinuria (≥300 mg/dl).

### Necropsy and tissue selection for light microscopy

Mice were anesthetized at the end of the study with IP injections at 250 mg/kg bw of sodium pentobarbital. A midline laparotomy was performed, and approximately 0.8 ml of blood was drawn from the inferior vena cava and collected in heparinized tubes (MiniCollect Capillary Blood Collection Tube #450477, Greiner Bio-One, Monroe, NC). Animals were euthanized by exsanguination through the inferior vena cava. After death, the trachea was exposed, cannulated, and the heart and lungs were excised en bloc. A volume of 0.7 ml sterile saline was instilled via the tracheal cannula and then withdrawn to recover bronchoalveolar lavage fluid (BALF). A second intratracheal saline lavage was performed, and the collected BALF was combined with the first sample. Organs selected for light microscopy and digital image analysis were harvested at necropsy and fixed in 10% neutral buffered formalin for at least 48 hours. The left kidney, spleen, and brain were fixed by immersion, while the left lung lobe was fixed by fluid inflation via the trachea at 30 cm gravitational pressure as previously described [57].

After fixation, cross-sectional (transverse/coronal) tissue blocks were cut from the left kidney (1-2 blocks covering the entire organ), left lung lobe (4 blocks covering the entire lobe), spleen (2 mid-blocks from the body of the organ), and brain (1 block with hippocampus, at the level of the inner ear). Tissue blocks were paraffin-embedded and sectioned at 5 microns. Tissue sections were stained histochemically with Hematoxylin and Eosin (H&E, all organs) or Periodic Acid Schiff (PAS, kidney) and immunohistochemically stained for CD3^+^ and CD45R^+^ T and B lymphoid cells, respectively (lung and kidney), IBA-1^+^ monocytes/macrophages (lung and kidney), and IBA-1^+^ microglia in the hippocampus (brain). Details for immunohistochemistry and specific antibodies are provided in the Supplemental Materials.

### Quantitative and semi-quantitative image analysis of histopathology

Digital images of histological sections were obtained with an Olympus VS200 slide scanner with ASW software (V.3.2.1) at 20x magnification (Evident Scientific, Waltham, MA). Digitally scanned images were viewed with Olympus OlyVIA software (V.2.8) and whole-slide image analyses were conducted using QuPath image analysis software (V.0.5.1) [68]. Regions of interest (ROIs) were annotated prior to each quantitative analysis (Lung, whole lung lobe sections; Kidney, renal cortex in sections). Abundance of immunohistochemical stains (density of IBA-1^+^, CD3^+^, CD45R^+^) was graphically expressed as the percent of ROI area. Thresholds were created to model staining positivity per tissue type. Identical settings per stain were used to assess positivity among experimental groups.

Semi-quantitative image analysis of histopathologic severity was conducted for specific lupus-related lesions: membranous glomerulonephritis, renal tubular regeneration, accumulated protein in renal tubular lumens, ectopic lymphoid structures (ELS, CD3^+^ T cells and CD45R^+^ B cells) in the kidney, white pulp hyperplasia in the spleen, and microgliosis in the hippocampus of the brain (IBA-1^+^ microglia). A board-certified veterinary pathologist (JRH) examined all digitally scanned images of routine H&E- and immunohistochemically stained tissue sections for cSiO_2_-triggered lupus histopathology using unbiased blinded assessment (no previous knowledge of individual animal exposure or treatment). Lesion severity scores were recorded in a manner that is consistent with state-of-the-art guidelines for semi-quantitative pathology assessment [69]. Criteria for ordinal scoring were defined by percent of total tissue affected: 0 (no lesions) = 0% of total tissue; 1 (minimal) = less than 5%; 2 (mild) = 6 to 10%; 3 (moderate) = 11 to 50%; 4 (marked) = 51 to 75%; 5 (severe) = greater than 76%.

### Red blood cell (RBC) fatty acid analysis

RBC fatty acid composition was analyzed using gas chromatography (GC) with flame ionization detection by OmegaQuant Analytics (Sioux Falls, SD) as described previously [63]. Briefly, after isolating RBCs from heparinized blood, fatty acid methyl esters (FAMEs) were prepared by heating the RBCs with 14% boron trifluoride-ethanol and hexane at 100°C for 10 minutes. The FAMEs were then extracted with hexane and analyzed on a Shimadzu GC-2010 chromatograph equipped with an SP-2560 100-m capillary column. Fatty acids were identified by comparison to a commercial standard (GLC OQ-A, NuCheck Prep). A total of 24 fatty acids were quantified and expressed as a percentage of the total identified fatty acids, including saturated, monounsaturated (cis and trans), and polyunsaturated (omega-3 and omega-6) species.

### Autoantibody (AAb) measurement

IgG AAbs targeting silica-killed cell (SKC) autoantigens (AAgs) and dsDNA were measured using an indirect ELISA based on a previously described method [70]. To prepare SKC AAgs, fetal liver-derived alveolar-like macrophages (FLAMs) from lupus-prone NZBWF1 mice were exposed to 50 µg/ml cSiO_2_ for 20 hours to induce cell death. The supernatant was then collected and stored at - 20°C until use. Poly-L-lysine-coated 96-well microplates were incubated with a blocking buffer (PBS, 2% [w/v] BSA, 0.05% [v/v] Tween 20), then coated with 50 µl per well of either SKC AAg diluted 1:20 or calf thymus dsDNA diluted to 2.5 µg/ml, both prepared in ELISA buffer (PBS, 0.1% [w/v] BSA, 0.05% [v/v] Tween 20). Next, diluted BALF (1:1,000) or plasma (1:5,000 and 1:10,000) was added to separate wells. A standard curve was generated using two-fold serial dilutions of a mouse anti-dsDNA antibody, with the highest concentration assigned a value of 2,000 arbitrary units (U). Bound AAbs were detected using HRP-conjugated goat anti-mouse IgG Fc antibody and K-Blue® Advanced Plus TMB Substrate, with absorbance measured at 650 nm.

### Blood urea nitrogen (BUN) analysis

BUN was measured using the Ray Biotech Urea Nitrogen Detection Assay Kit (MA-BUN-5) per manufacturer’s directions. Briefly, plasma samples were diluted 1:20, loaded onto a microplate, and then incubated with an enzyme solution of urease and NADH. After a brief shaking of the plate, absorbance was measured (340 nm) twice, with one minute between readings. The difference in absorbance between the two readings correlates with BUN concentration. BUN levels were determined using a standard curve.

### Bronchoalveolar lavage fluid (BALF) cell quantitation and identification

Total cells in BALF were counted using a hemocytometer. Cytological slides were prepared by centrifuging 150 μl of BALF for 10 minutes at 40 × *g*, air-drying, and staining with the Easy III Step Stain Kit (#ES902, Azer Scientific, Morgantown, PA). At least 150 cells were counted and differentially identified as monocytes/macrophages, lymphocytes, or neutrophils based on morphological criteria.

### Statistical analysis

All treatment groups began with 8 mice. All statistical analyses were conducted using GraphPad Prism 10 (V.10.6.1, Dotmatics, Boston, MA), with a p-value < 0.05 considered significant. Parametric two-tailed t tests (with and without Welch’s correction) or nonparametric Mann-Whitney tests were used to compare the continuous quantitative data and the ordinal histopathological severity scores between the cSiO_2_/VEH/CON (negative control) and cSiO_2_/CYC/CON (positive control) groups, respectively. When comparing the continuous quantitative data across the treatment groups (cSiO_2_/CYC/CON, cSiO_2_/CYC/DHA, cSiO_2_/CYC/PDN, cSiO_2_/CYC/DHA+PDN), several statistical tests were performed: (i) parametric one-way ANOVAs along with the Brown-Forsythe test of equal variances, (ii) parametric Brown-Forsythe and Welch one-way ANOVAs, and (iii) Dunnett’s and Tukey’s multiple comparisons tests. When comparing the semi-quantitative ordinal severity score data of the treatment groups, the nonparametric Kruskal-Wallis and Dunn’s multiple comparisons tests were used.

## Results

### Dietary DHA remodels RBC membrane fatty acid composition

The dietary DHA intervention was confirmed to induce systemic lipidomic changes by assessing the fatty acid composition of RBC phospholipids, which serves as a biomarker of overall tissue lipid status (Figure 2; Table 3) [71]. Mice receiving isocaloric DHA-supplemented diets (cSiO₂/CYC/DHA and cSiO₂/CYC/DHA+PDN groups) exhibited significant increases (up to 10- to 12-fold) in DHA and other omega-3 fatty acids, including EPA and omega-3 docosapentaenoic acid (DPA). This increase was accompanied by a marked decrease in total omega-6 fatty acids (up to 50 to 70%), such as ARA, omega-6 DPA, and docosatetraenoic acid, with the exception of linoleic acid (LA), which was significantly elevated. Notably, RBCs from mice treated with DHA and PDN contained higher levels of DHA, EPA, and total omega-3s than those from mice receiving DHA alone, along with lower levels of ARA and total omega-6 fatty acids. In contrast, PDN treatment alone did not alter fatty acid profiles, indicating that the observed lipidomic changes were specifically attributable to dietary DHA intake.

**Figure 2.**
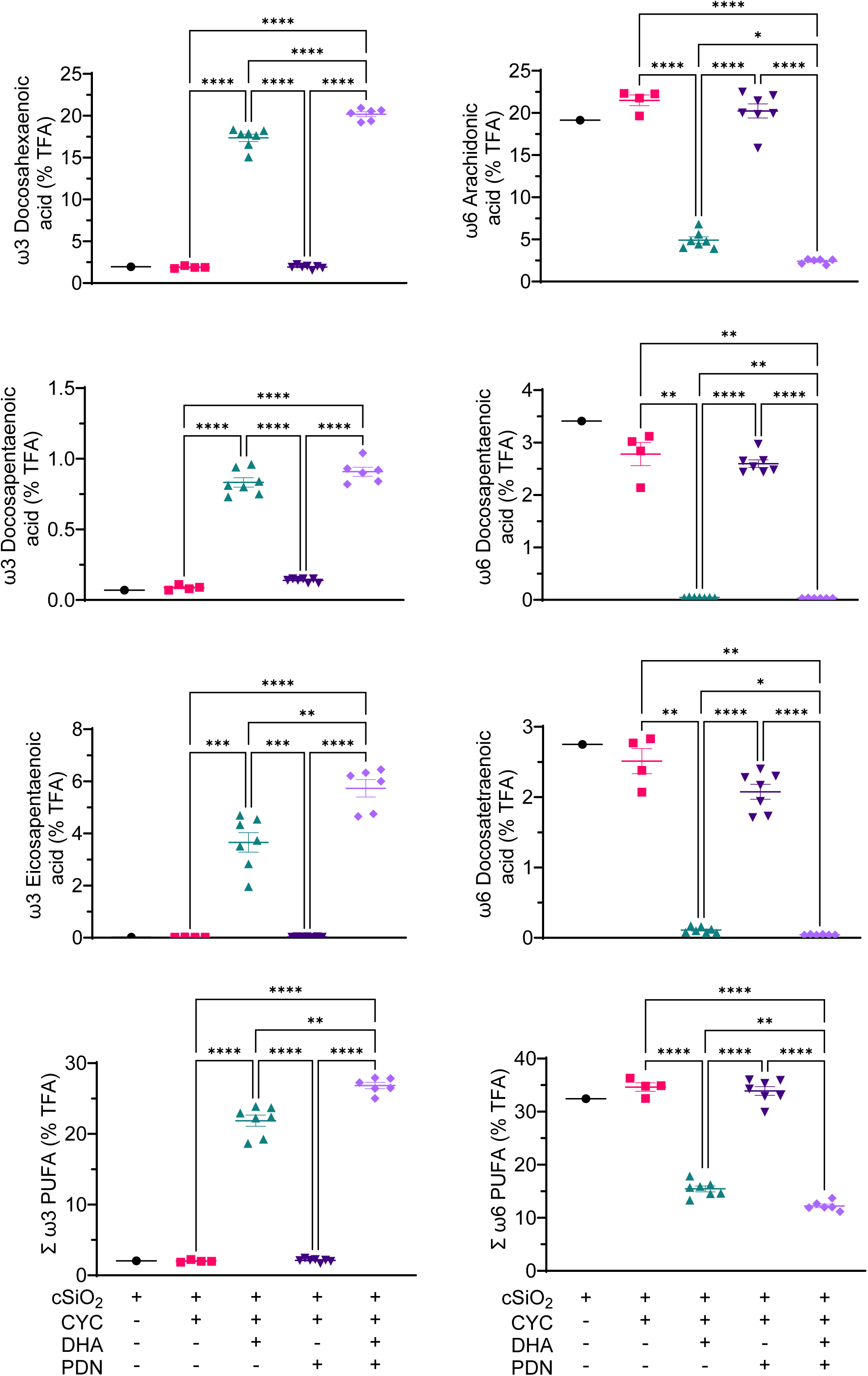
Consumption of DHA and DHA+PDN, but not PDN alone, skews cellular phospholipid profiles in erythrocytes from ω-6 PUFAs to ω-3 PUFAs. Data (mean+/-SEM) are expressed as percent of total fatty acids (TFA). *, **, ***, and **** indicate significant differences at the p<0.05, p<0.01, p<0.001, and p<0.0001 levels, respectively.

**Table 3.** Effects of experimental diets on fatty acids in red blood cells (RBCs).

| Chemical<br>Structure | Experimental Group |  |  |  |
| --- | --- | --- | --- | --- |
|  | Percent (%) of Total Fatty Acids |  |  |  |
|  | cSiO<sub>2</sub>/CYC/CON | cSiO<sub>2</sub>/CYC/DHA | cSiO<sub>2</sub>/CYC/PDN | cSiO<sub>2</sub>/CYC/DHA+PDN |
| C14:0 | 0.11 ± 0.02 | 0.26 ± 0.07** | 0.13 ± 0.03 | 0.30 ± 0.08** |
| C16:0 | 25.76 ± 1.6 | 29.92 ± 0.81**** | 29.74 ± 1.11 | 25.76 ± 1.9**** |
| C16:1ω7t | 0.05 ± 0.008 | 0.05 ± 0.005 | 0.05 ± 0.008 | 0.05 ± 0.01 |
| C16:1ω7 | 0.56 ± 0.07 | 0.67 ± 0.20 | 0.95 ± 0.22* | 1.03 ± 0.20** |
| C18:0 | 12.17 ± 1.5 | 10.83 ± 1.09 | 10.96 ± 1.53 | 10.75 ± 0.89 |
| C18:1t | 0.12 ± 0.03 | 0.10 ± 0.02 | 0.11 ± 0.01 | 0.08 ± 0.02* |
| C18:1ω9 | 23.33 ± 3.56 | 19.74 ± 1.35 | 24.63 ± 3.37 | 17.82 ± 1.26** |
| C18:2ω6t | 0.10 ± 0.02 | 0.09 ± 0.02 | 0.14 ± 0.02** | 0.09 ± 0.02 |
| C18:2ω6 | 6.51 ± 1.44 | 9.14 ± 0.63**** | 7.49 ± 0.72 | 8.92 ± 0.89*** |
| C20:0 | 0.11 ± 0.02 | 0.10 ± 0.02 | 0.10 ± 0.01 | 0.12 ± 0.03 |
| C18:3ω6 | 0.05 ± 0.02 | 0.03 ± 0.01* | 0.07 ± 0.01** | 0.03 ± 0.009* |
| C20:1ω9 | 0.46 ± 0.05 | 0.24 ± 0.02**** | 0.38 ± 0.08 | 0.19 ± 0.04**** |
| C18:3ω3 | 0.01 ± 0 | 0.01 ± 0.006 | 0.01 ± 0.007 | 0.02 ± 0.006 |
| C20:2ω6 | 0.19 ± 0.04 | 0.16 ± 0.02 | 0.16 ± 0.02 | 0.11 ± 0.02**** |
| C22:0 | 0.12 ± 0.03 | 0.14 ± 0.04 | 0.12 ± 0.02 | 0.14 ± 0.08 |
| C20:3ω6 | 1.00 ± 0.10 | 0.97 ± 0.13 | 1.14 ± 0.14 | 0.61 ± 0.13*** |
| C20:4ω6 | 21.49 ± 2 | 4.91 ± 0.94**** | 20.23 ± 2.05 | 2.40 ± 0.31**** |
| C24:0 | 0.22 ± 0.06 | 0.28 ± 0.07 | 0.23 ± 0.03 | 0.38 ± 0.06*** |
| C20:5ω3 | 0.02 ± 0.008 | 3.66 ± 0.92**** | 0.03 ± 0.01 | 5.73 ± 0.85**** |
| C24:1ω9 | 0.37 ± 0.08 | 0.35 ± 0.08 | 0.33 ± 0.04 | 0.35 ± 0.05 |
| C22:4ω6 | 2.51 ± 0.57 | 0.11 ± 0.04** | 2.08 ± 0.26 | 0.04 ± 0.005** |
| C22:5ω6 | 2.78 ± 0.70 | 0.05 ± 0.007** | 2.60 ± 0.17 | 0.03 ± 0.004** |
| C22:5ω3 | 0.09 ± 0.03 | 0.83 ± 0.08**** | 0.14 ± 0.01 | 0.91 ± 0.08**** |
| C22:6ω3 | 1.89 ± 0.23 | 17.35 ± 1.1**** | 1.92 ± 0.20 | 20.17 ± 0.75**** |
| DHA + EPA | 1.91 ± 0.24 | 21.01 ± 1.87**** | 1.95 ± 0.21 | 25.9 ± 1.2**** |
| CX:0 | 38.50 ± 1.65 | 41.54 ± 0.77** | 37.54 ± 1.59 | 41.42 ± 0.58** |
| CX:1 | 24.88 ± 3.51 | 21.15 ± 1.42 | 26.46 ± 3.61 | 19.52 ± 1.4** |
| CX:≥2ω3 | 2.01 ± 0.26 | 21.86 ± 1.93**** | 2.10 ± 0.21 | 26.83 ± 1.16**** |
| CX:≥2ω6 | 34.62 ± 2.51 | 15.45 ± 1.34**** | 33.90 ± 2.01 | 12.24 ± 0.9**** |

Fatty acid profiles of RBCs were determined by OmegaQuant Analytics as described in Methods. Data are expressed as mean ± SEM. Symbols: \*, \*\*, \*\*\*, and \*\*\*\* indicate significant differences at the p<0.05, p<0.01, p<0.001, and p<0.0001 levels, respectively, when compared to the cSiO<sub>2</sub>/CYC/CON group.

### DHA and PDN co-therapy after CYC induction prevents proteinuria relapse and death

Consistent with previous studies in NZBWF1 mice [54,56,61], weekly intranasal cSiO_2_ instillations for 4 weeks in control-fed mice resulted in two main clinical outcomes: proteinuria (Figures 3A-C) and shortened lifespan, with a median survival of 29.5 weeks (Figure 3D). CYC treatment alone reduced proteinuria, but relapses happened after stopping, as evidenced by increased proteinuria at week 33 (Figure 3A), and increased median survival to 35 weeks (Figure 3D). Maintenance therapies with either DHA or DHA+PDN delayed relapse, whereas PDN alone did not. The mean percentage of proteinuric mice (Figure 3B) and mean proteinuria levels (Figure 3C) from weeks 21 to 35 were significantly lower in the CYC/DHA and CYC/DHA+PDN groups than in the CYC-only or CYC/PDN groups. Maintenance treatments also conferred a survival advantage, with median survival times for both monotherapies and combination therapy exceeding 35 weeks (Figure 3D).

**Figure 3.**
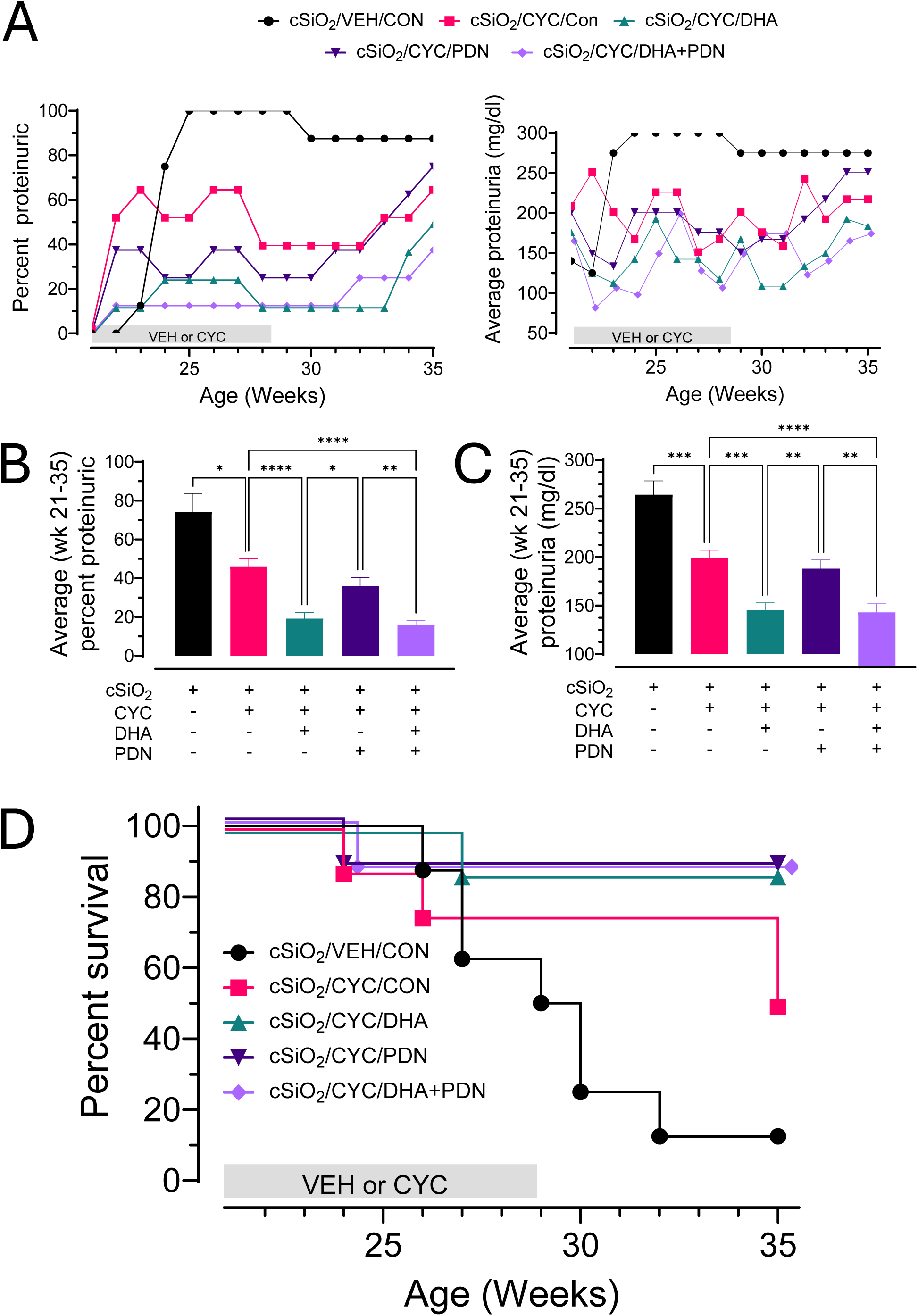
DHA and PDN monotherapies and co-therapy influence the duration of CYC-induced remission of proteinuria and mortality in the SALN model over the 14-week experimental period. (A) Left panel, percent of mice with significant proteinuria (≥300 mg/dl); right panel, average proteinuria. (B) Average percent of mice with proteinuria (≥300 mg/dl) from weeks 21 to 35. (C) Average proteinuria from weeks 21 to 35. (D) Percent survival as determined using clinical scores indicative of a moribund state as described in Methods. Abbreviations: VEH, vehicle; CON, control diet. Symbols: *, **, ***, and **** indicate significant differences at the p<0.05, p<0.01, p<0.001, and p<0.0001 levels, respectively.

### Combining DHA and PDN maintenance therapy preserves renal structure and reduces macrophage infiltration

Kidneys from cSiO₂/VEH/CON mice exhibited severe membranous glomerular nephritis, characterized by glomerular enlargement, extensive accumulation of PAS^+^ glomerular membranous tissue, and proteinaceous casts in the renal tubules (Figure 4B). These features were absent in age-equivalent naïve mice from a prior study [61] and used here as a standard reference control (Figure 4A). Conversely, CYC treatment reduced glomerulonephritis (Figure 4C), and this improvement was further enhanced by DHA (Figure 4D) and PDN (Figure 4E). The group receiving combination therapy (cSiO₂/CYC/DHA+PDN) was the most protected from these pathological changes and displayed nearly normal renal structure (Figure 4F). Semi-quantitative scoring confirmed that the combination therapy decreased the LN severity score (p<0.01) (Figure 4G), tubular protein score (p=0.11) (Figure 4H), and tubular regeneration score (p<0.05) (Figure 4I) compared to CYC treatment alone.

**Figure 4.**
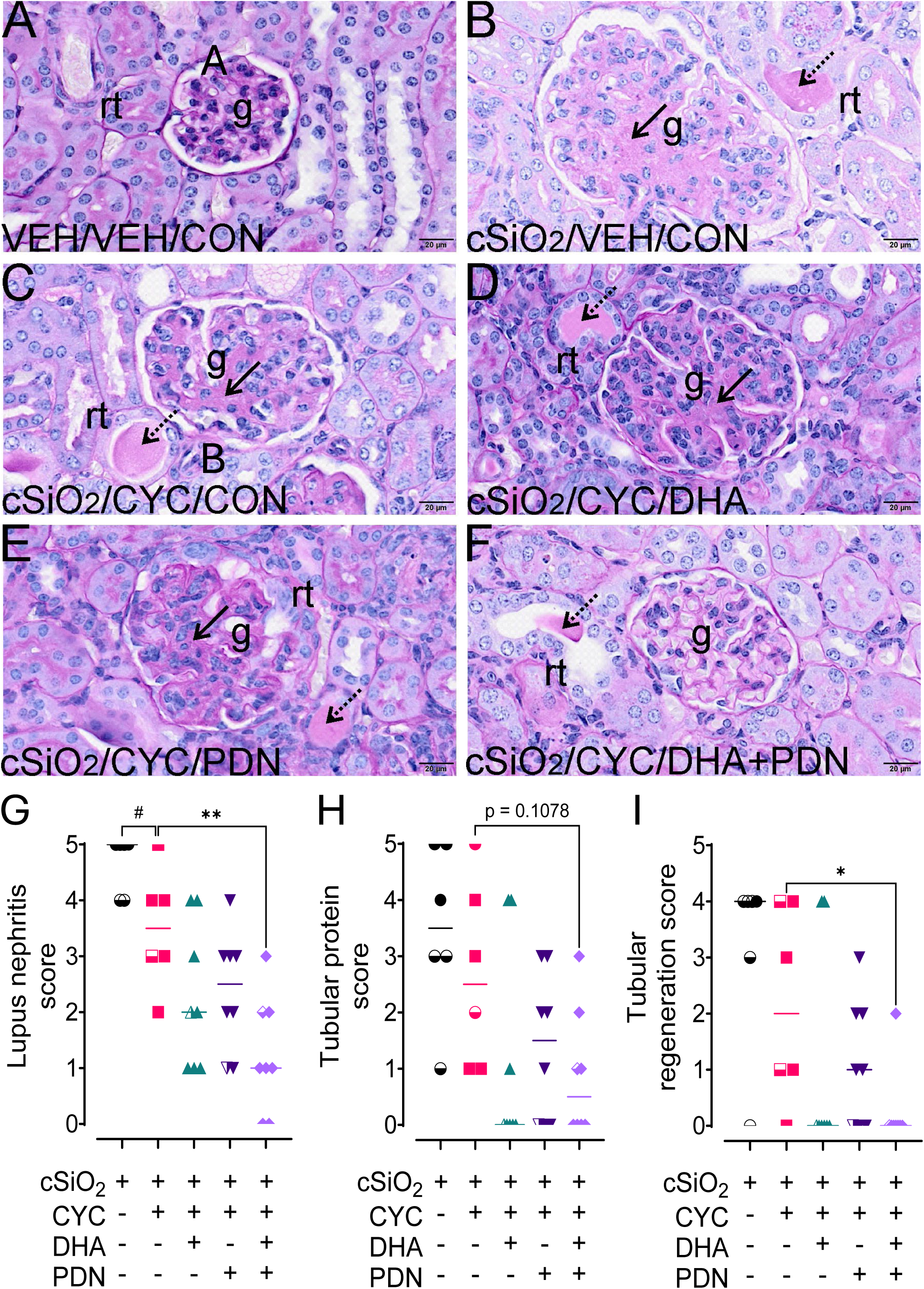
DHA and PDN monotherapies and co-therapy prolong CYC-induced diminution of cSiO_2_-accelerated lupus nephritis. (A-F) Light photomicrographs of Periodic Acid Schiff-stained (PAS) glomeruli (g) and adjacent cortical renal tubules (rt). (B) Membranous glomerular nephritis: dramatic glomerular enlargement with marked thickening of PAS^+^ membranous tissue (arrow) and rt with luminal proteinaceous fluid (stippled arrow) in the kidney from the cSiO_2_/VEH/CON mouse compared to (A) a representative section from a 34-week-old NZBWF1 mouse not treated with cSiO_2_ or drugs obtained from a previous study [61]. These renal lesions are markedly attenuated in the kidney from the (F) cSiO_2_/CYC/DHA+PDN mouse. Histopathological lesions were scored by pathologist as described in Methods for (G) lupus nephritis, (H) tubular protein, and (I) tubular regeneration. Symbols: #, *, and ** denote statistically significant differences at the p<0.1, p<0.05, and p<0.01 levels, respectively. Solid symbols indicate mice that were taken at the final necropsy at week 35; half-open, half-closed symbols denote moribund mice sacrificed prior to final necropsy.

IHC staining for IBA-1, a well-known marker for recruited inflammatory monocytes/macrophages [72], showed that, consistent with glomerulonephritis, cSiO₂ exposure caused significant infiltration of monocytes/macrophages into interstitial spaces surrounding renal tubules and glomeruli, in ELS, and some glomeruli (Figures 5A, 5B). While modestly reduced by CYC with DHA or PDN monotherapies (Figures 5D, 5E, 5G), IBA-1^+^ monocyte/macrophage infiltration was most robustly reduced by co-therapy with CYC/DHA+PDN (Figures 5F, 5G). Accordingly, DHA+PDN co-treatment was the most effective of the three maintenance therapies in reducing local renal inflammation and preserving kidney structure.

**Figure 5.**
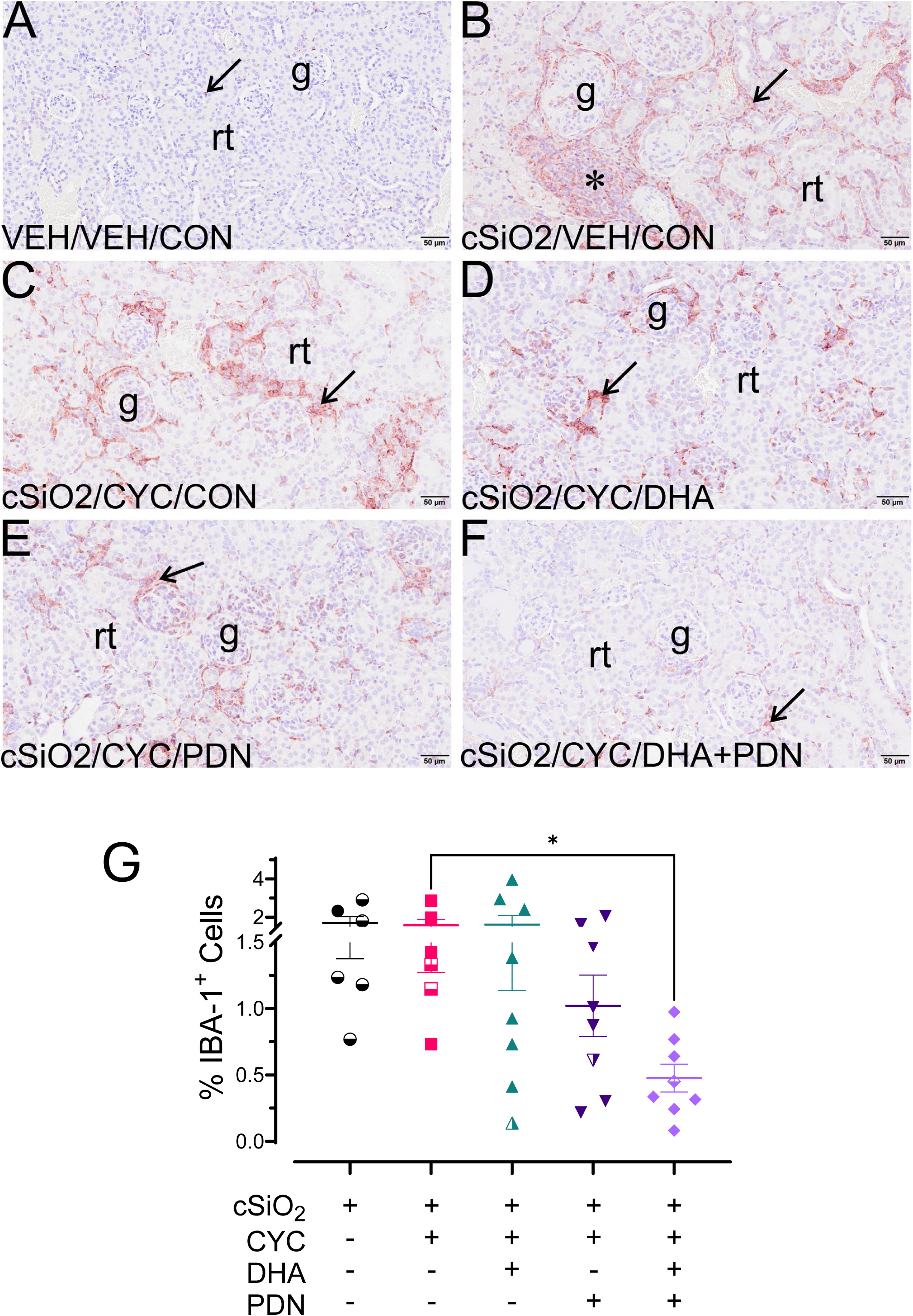
DHA and PDN co-therapy extends CYC-induced suppression of cSiO_2_-induced infiltration of inflammatory IBA-1^+^ monocyte-derived macrophages in the kidney. (A-F) Representative light photomicrographs of renal cortex immunohistochemically stained for IBA-1^+^ monocytes infiltrating peri-tubular and glomerular interstitial tissue (arrows). Panel A is a representative section from a 34-week-old NZBWF1 mouse not treated with cSiO_2_ or drugs obtained from a previous study [61]. Letters: g, glomeruli; rt, renal tubules. (G) IBA-1^+^ areas were quantitatively scored as described in Methods. Symbol: * denotes a statistically significant difference at the p<0.05 level. Solid symbols indicate mice that were taken at the final necropsy at week 35; half-open, half-closed symbols denote moribund mice sacrificed prior to final necropsy.

### DHA and PDN co-therapy prevents the formation of renal ELS

A key pathogenic feature of progressive LN is the formation of renal ELS [73–75]. The renal cortices of cSiO_2_-triggered mice that did not receive treatment contained large, organized ELS composed of co-localized aggregations of IBA-1⁺ macrophages, CD3⁺ T cells, and CD45R⁺ B cells (Figure 6A, C, E, G). Maintenance therapy with CYC/DHA+PDN effectively prevented this process, as kidneys from this group were mostly free of dense lymphoid cell aggregate infiltrates and lacked organized ELS architecture (Figure 6B, D, F, H). The kidney ELS score was lower (p<0.05) in the CYC/DHA+PDN co-therapy group compared to the CYC-only and CYC/DHA groups, while the PDN monotherapy was less effective (Figure 6I).

**Figure 6.**
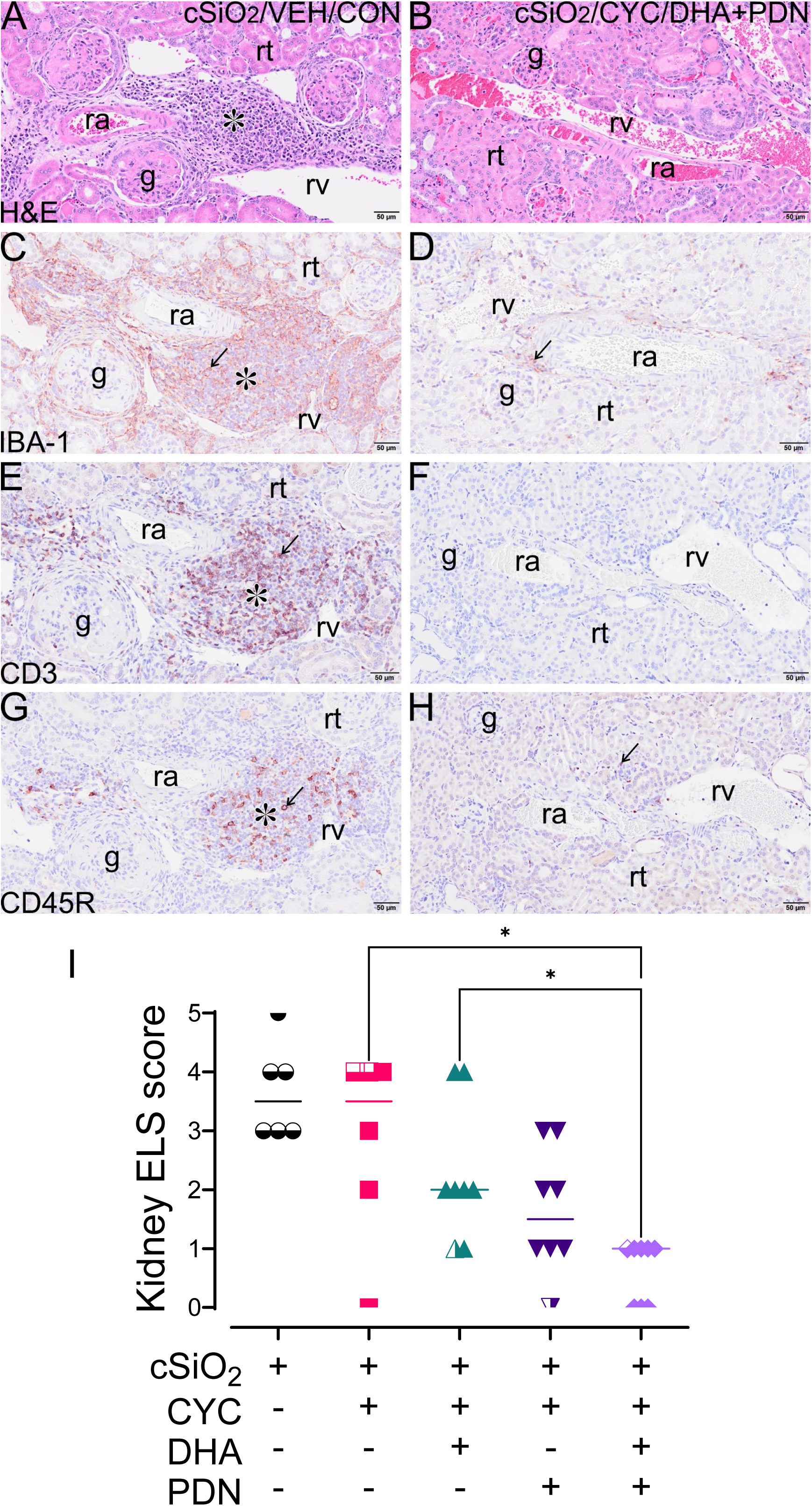
DHA and PDN co-therapy prolongs CYC-induced attenuation of cSiO_2_-accelerated ectopic lymphoid structure (ELS) development in the kidney. Representative light photomicrographs of (A, B) hematoxylin and eosin (H&E)-stained renal cortex tissue and immunohistochemically stained for (C, D) IBA-1^+^, (E, F) CD3^+^, or (G, H) CD45R^+^ cells in renal cortex tissue from a (A, C, E, G) nontreated cSiO_2_/VEH/CON mouse and a (B, D, F, H) co-treated cSiO_2_/CYC/DHA+PDN mouse. cSiO_2_-induced renal influx of IBA-1^+^ monocytes co-localized with CD3^+^ and CD45R^+^ lymphoid cells (T and B cells, respectively) was markedly attenuated in co-treated mouse compared to nontreated mouse. (I) ELS development was semi-quantitatively scored for severity by pathologist as described in Methods. Symbol: * indicates a significant difference at the p<0.05 level. Solid symbols indicate mice that were taken at the final necropsy at week 35; half-open, half-closed symbols denote moribund mice sacrificed prior to final necropsy.

### Protective effects of DHA and PDN monotherapy and co-therapy extend to the lungs

In NZBWF1 mice, age-related pulmonary perivascular and peribronchiolar lymphoid hyperplasia spontaneously occurs at 8 to 12 months of age, during which polymorphic lymphoid infiltrates, evocative of ELS, become dominated by CD4^+^ T cells, while B cells and antigen-presenting cells segregate into a peripheral cuff surrounding the T cell–rich area [76,77]. This occurs much earlier in the SALN model, where the lung is the hub for early cellular events that promote cSiO_2_-induced autoimmunity and the progression of lupus [78]. Cellular analysis of BALF demonstrated that DHA and PDN monotherapies, as well as combination therapy with CYC, reduced total inflammatory cells, monocytes/macrophages, and lymphocytes compared to CYC alone (Figure 7A). Furthermore, combined DHA+PDN treatment exhibited a modest trend toward reduced local production of IgG autoantibodies against dsDNA and cSiO_2_-killed cells (Figure 7B).

**Figure 7.**
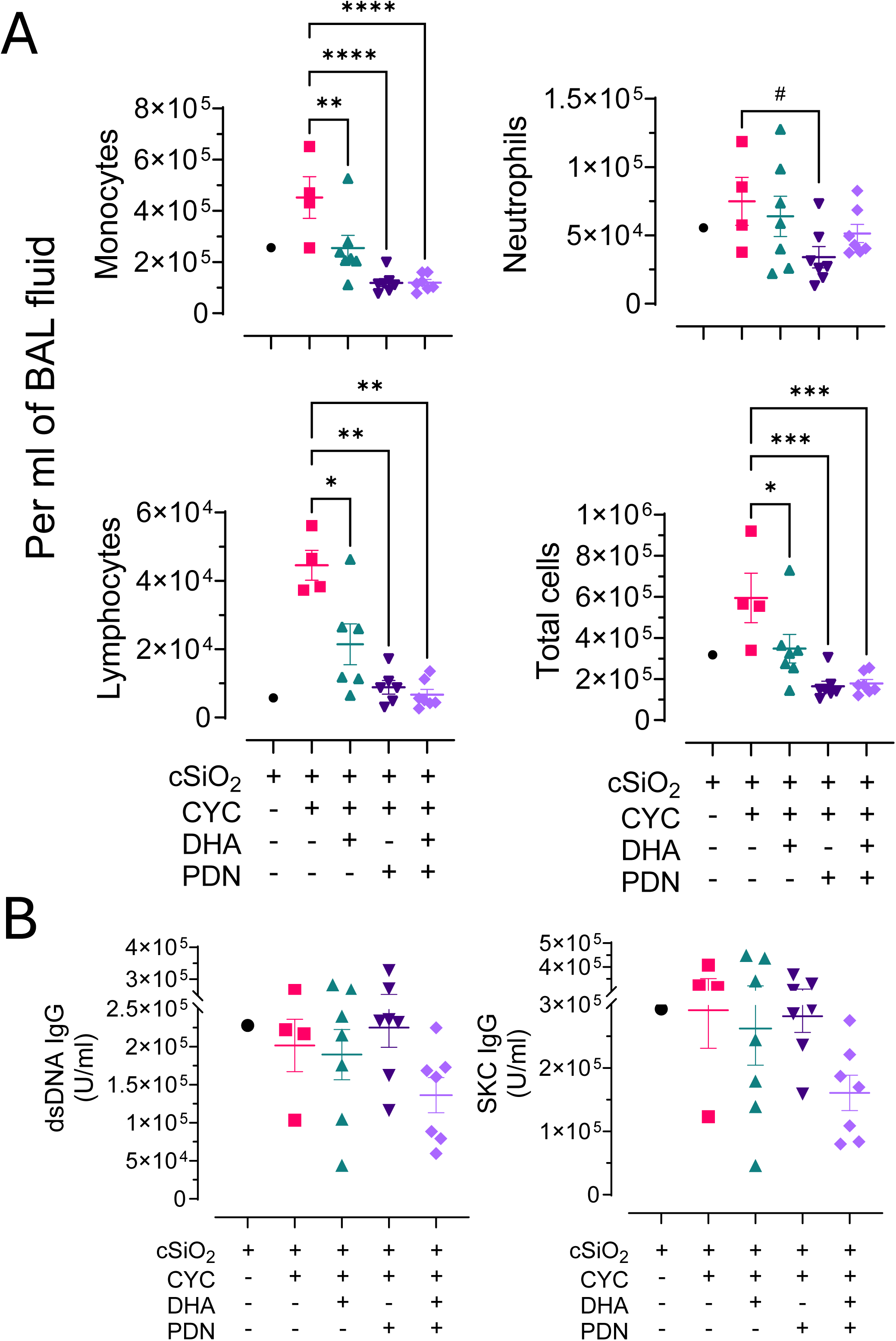
Influence of DHA and PDN monotherapies and co-therapy on cSiO_2_-induced cellularity and AAbs in bronchoalveolar lavage fluid (BALF). (A) Monocyte, neutrophil, lymphocyte, and total cell accumulation are suppressed by DHA and PDN alone or as co-treatment. Symbols: #, *, **, ***, and **** indicate significant differences at the p<0.1, p<0.05, p<0.01, p<0.001, and p<0.0001 levels, respectively. (B) IgG AAbs specific for dsDNA and silica-killed cells (SKC) of CYC-treated mice instilled with cSiO_2_ are modestly suppressed by DHA and PDN co-treatment.

Histological examination of lung tissue showed that, as previously reported [54,55], strong perivascular ELS appeared in cSiO_2_-treated mice (Figures 8C, D) and were absent in naïve mice (Figures 8A, B). Consistent with decreased cellularity in BALF, ELS were markedly reduced in the lungs of the CYC/DHA+PDN group (Figures 8K, L), while effects of monotherapies were more modest (Figures 8G-J). Semi-quantitative scoring confirmed that the DHA+PDN combination maintenance therapy significantly lowered lung ELS scores (p<0.001 and p<0.05) compared to CYC-only and CYC/PDN, respectively (Figure 9A) and modestly decreased the percentage of CD3^+^ T cells (Figure 9C), while both CYC/PDN and CYC/DHA+PDN significantly decreased the percentage of CD45R^+^ B cells (p<0.05 and p<0.01) compared to CYC-only, respectively (Figure 9D).

**Figure 8.**
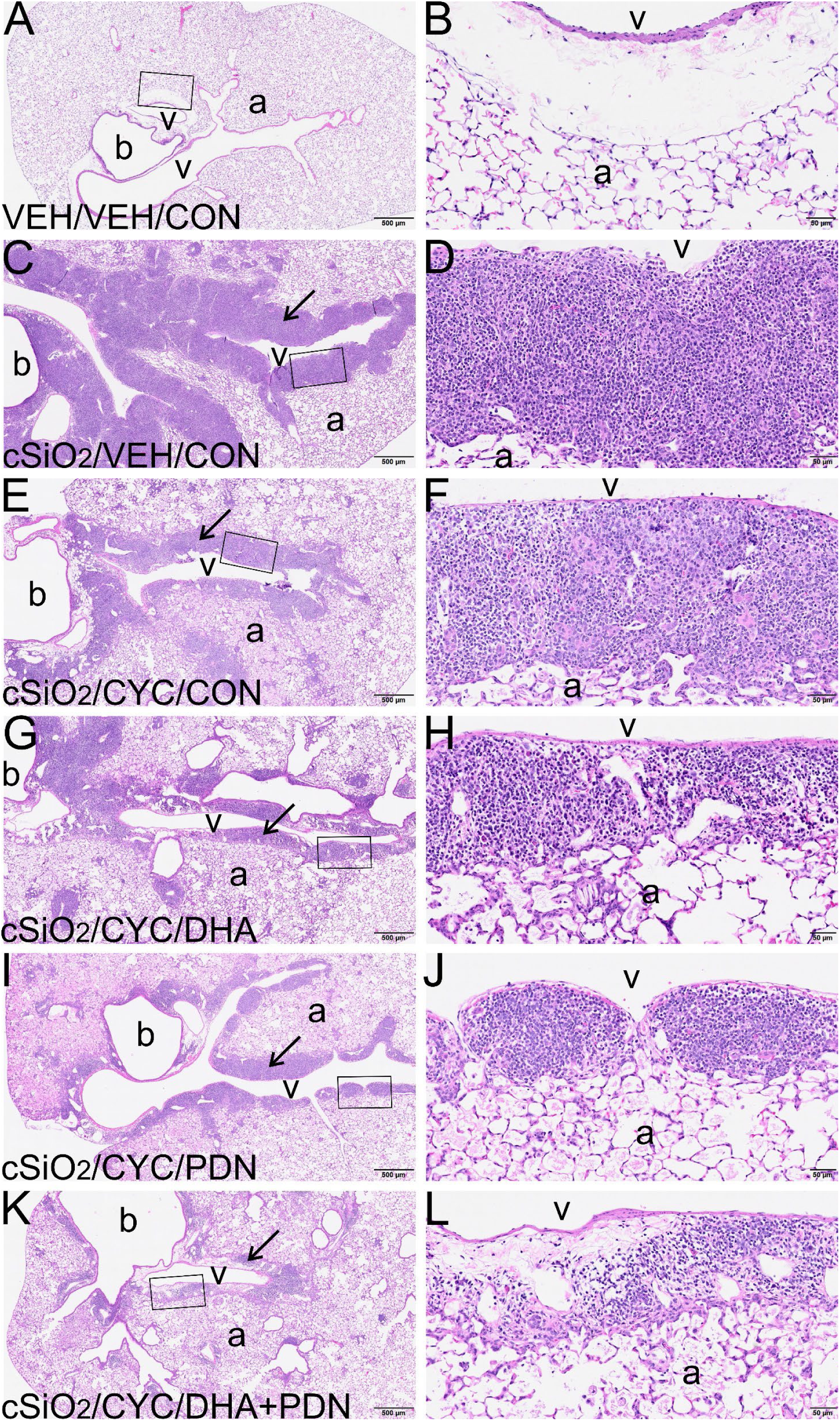
DHA and PDN monotherapies and co-therapy extend CYC-induced attenuation of cSiO_2_-accelerated ectopic lymphoid structure (ELS) development in the lung. (A-L) Representative light photomicrographs of H&E-stained pulmonary tissue from the left lung lobe. Panels A and B are a representative section from a 34-week-old NZBWF1 mouse not treated with cSiO_2_ or drugs obtained from a previous study [61]. Peri-vascular ELS (arrows) is markedly attenuated in lung from (K, L) co-therapy cSiO_2_/CYC/DHA+PDN mouse compared to (C, D) nontreated cSiO_2_/VEH/CON and (E, F) cSiO_2_/CYC/CON mice.

**Figure 9.**
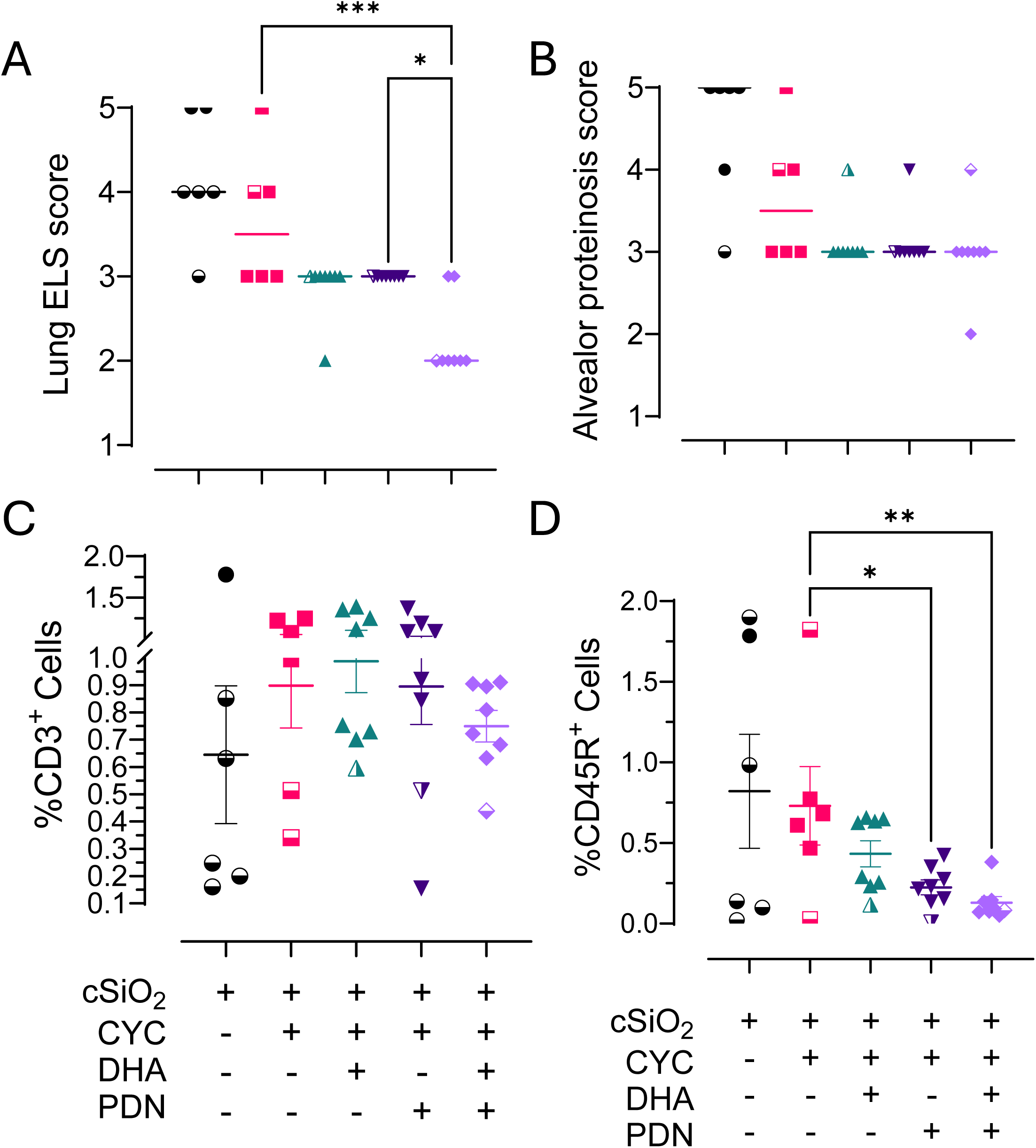
DHA and PDN monotherapies and co-therapy prolong CYC-induced reduction of cSiO_2_-accelerated ectopic lymphoid structure development in the lung. (A, B) ELS development and alveolar proteinosis were semi-quantitatively scored for severity by pathologist as described in Methods. (C, D) CD3^+^ and CD45R^+^ cell densities were quantitatively analyzed by digital morphometry as described in Methods. Symbols: *, **, and *** indicate significant differences at the p<0.05, p<0.01, and p<0.001, levels, respectively. Solid symbols indicate mice that were taken at the final necropsy at week 35; half-open, half-closed symbols denote moribund mice sacrificed prior to final necropsy.

### DHA+PDN co-therapy suppresses cSiO_2_-induced splenomegaly and systemic AAbs

The spleen is another potential source of systemic AAbs in NZBWF1 mice [79], and splenomegaly, a key feature of spontaneous lupus progression in these mice, is accelerated by cSiO_2_ treatment [64]. Therefore, to evaluate the effects of treatments on systemic autoimmunity, we compared spleen architecture and plasma AAbs across the experimental groups. cSiO_2_ caused significant white pulp hyperplasia in mice (Figures 10B, H), which was not evident in age-equivalent naïve NZBWF1 mice (Figure 10A). Lymphoid cell hyperplasia was not substantially affected by CYC alone (Figure 10C) or by CYC in combination with DHA or PDN alone (Figures 10D, E); however, it was suppressed in CYC-treated mice that received DHA+PDN (Figure 10F). For other indicators of splenomegaly, spleen weight and area, DHA and PDN monotherapies showed modest reductions, while DHA+PDN co-therapy showed marked decreases (Figures 10G, H, and I). Treatment effects on the spleen were consistent with plasma dsDNA and SKC AAb levels (Figure 10J). After stopping CYC, plasma IgG AAbs against dsDNA and SKC began to rise again within 4 to 6 weeks post-CYC treatment in the CYC-only and CYC/DHA groups. This increase was consistently suppressed in groups receiving PDN, either alone or with DHA. Overall, DHA potentiates PDN suppression of splenomegaly and systemic AAb formation resulting from cSiO_2_ instillation.

**Figure 10.**
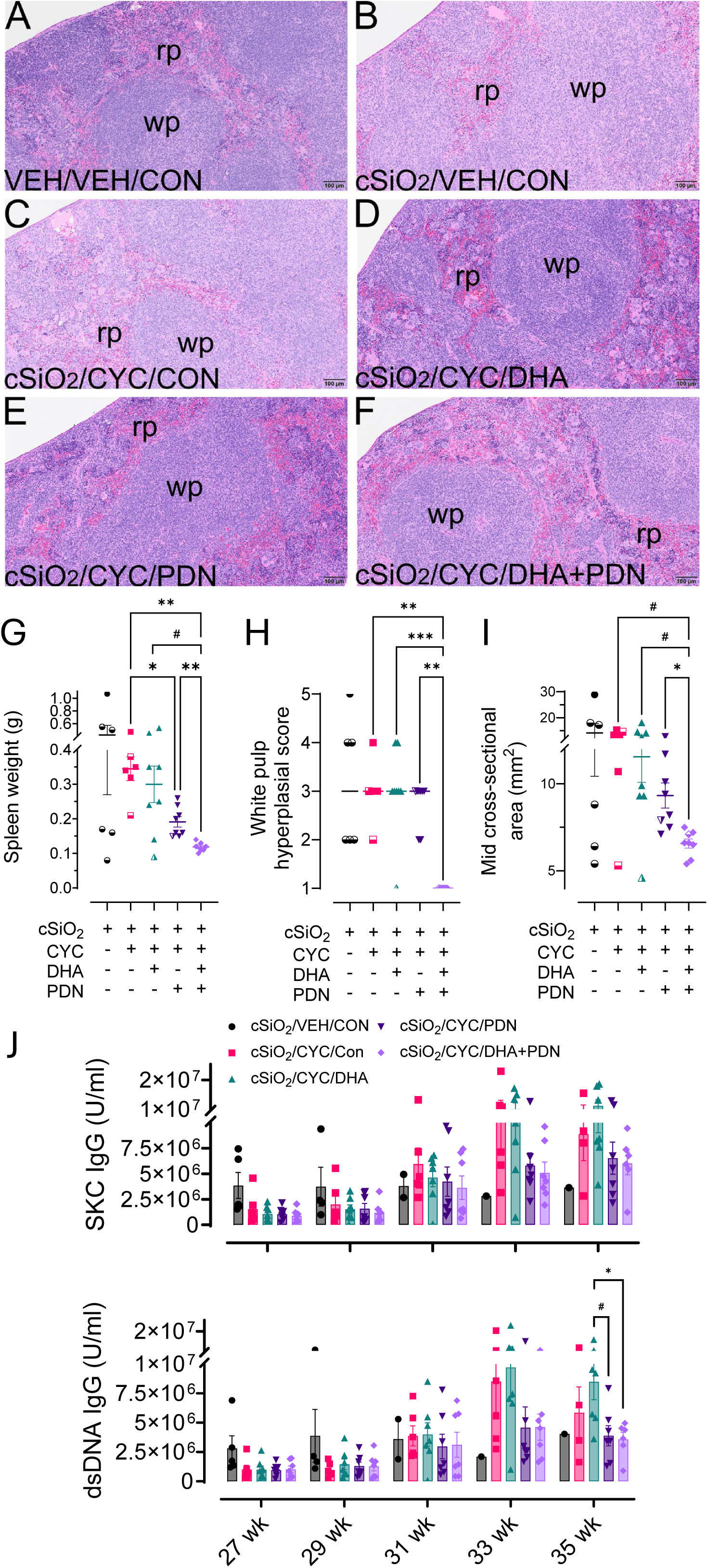
PDN alone or DHA and PDN co-therapy with CYC suppress cSiO_2_-accelerated splenomegaly. (A-F) Light photomicrographs of H&E-stained splenic tissue. (A) Reference section from a 34-week-old NZBWF1 mouse not treated with cSiO_2_ or drugs obtained from a previous study [61]. (B) Increased white pulp (wp) and reduced red pulp (rp) in splenic tissue from cSiO_2_/VEH/CON mouse that is markedly attenuated in (F) spleen from mouse receiving cSiO_2_/CYC/DHA+PDN treatment. Indicators of splenomegaly included (G) spleen weight, (H) white pulp hyperplasia (semi-quantitative scoring by pathologist as described in Methods), and (I) mid cross-sectional area of total spleen (quantitative analysis as described in Methods). (J) Influence of DHA and PDN monotherapies and co-therapy on cSiO_2_-induced AAbs in plasma. IgG AAbs specific for dsDNA and silica-killed cells (SKC) from CYC-treated mice instilled with cSiO_2_ are modestly suppressed by PDN treatment alone and by DHA and PDN co-treatment at 33 and 35 weeks. Symbols: #, *, **, and *** indicate significant differences at the p<0.1, p<0.05, p<0.01, and p<0.001 levels, respectively. Mice sacrificed prior to final necropsy are indicated by half-open, half-closed symbols, and solid symbols indicate mice that made it to final necropsy at week 35.

### DHA and PDN co-therapy attenuates microglial cell activation in the brain

NZBWF1 mice have been used in neuropsychiatric lupus (NPSLE) research, with IBA-1 positivity in the brain (i.e., microgliosis) and microglial activation serving as markers of central nervous system (CNS) inflammation [80–82]. In the hippocampus, cSiO₂/VEH/CON mice exhibited pronounced microgliosis, characterized by increased density and altered shape of IBA-1⁺ microglial cells (Figures 11A, B). This neuroinflammatory state was diminished in mice that received CYC, CYC/DHA, and CYC/PDN (Figures 11C, D, and E), and was maximally reduced in mice receiving CYC/DHA+PDN co-therapy, which displayed a more resting microglial appearance (Figure 11F). This was further reflected in markedly lower microgliosis scores for the DHA+PDN co-therapy group, suggesting that the benefits of this combination extend to reduced CNS-related inflammation (Figure 11G).

**Figure 11.**
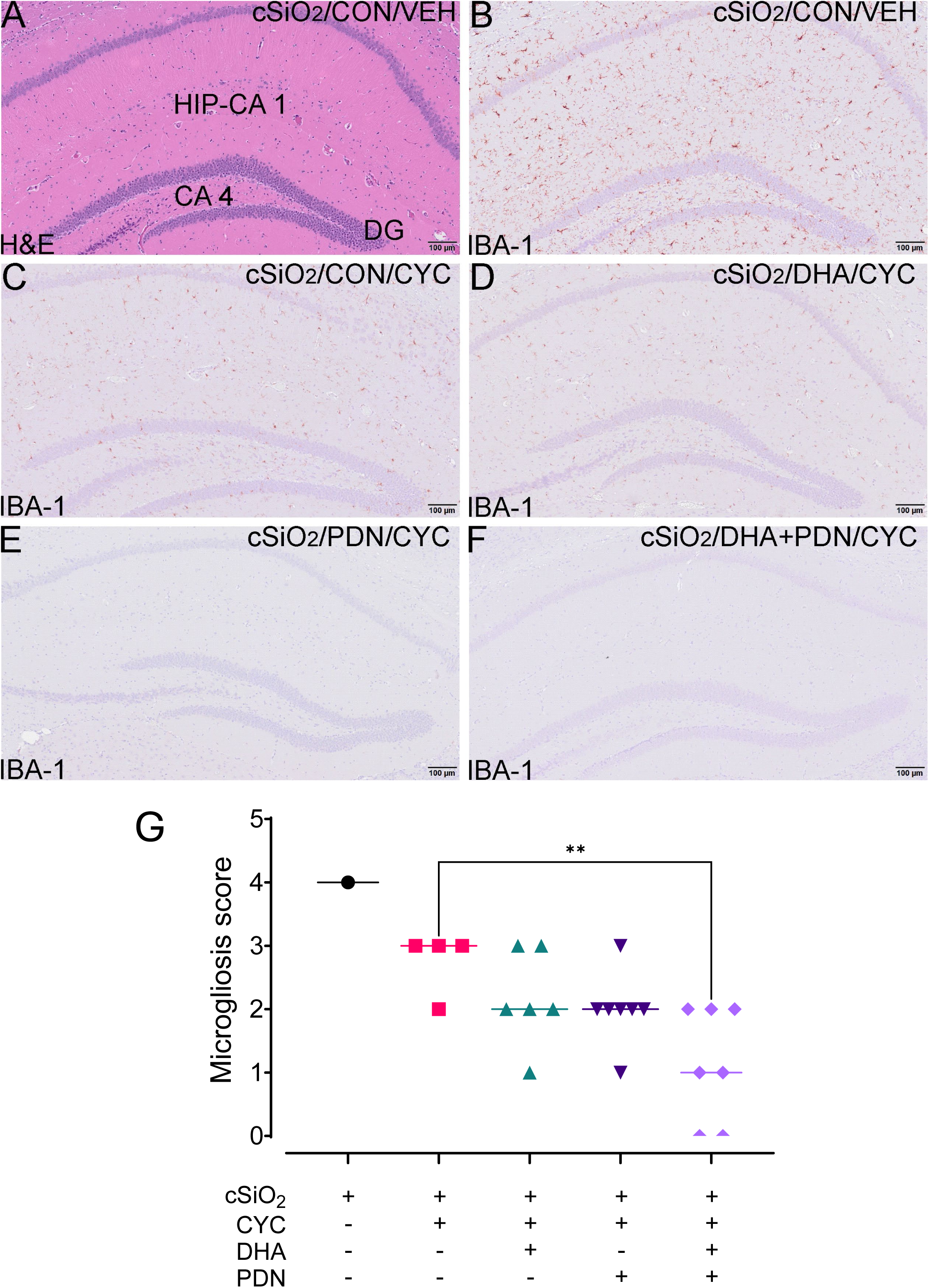
DHA and PDN monotherapies and co-therapy prolong CYC suppression of cSiO_2_-induced microgliosis. Representative photomicrographs of (A) H&E-stained cells and (B-F) immunohistochemically stained for IBA-1^+^ microglial cells in the hippocampus of the brain. Microgliosis in the hippocampus from the (B) nontreated cSiO_2_/VEH/CON mouse is markedly attenuated in (F) co-treated cSiO_2_/CYC/DHA+PDN mouse. (G) Semi-quantitative severity scoring of microgliosis (IBA-1^+^ cells) in the hippocampus as described in Methods. Brains were not taken from moribund mice at necropsy. Symbol: ** indicates a significant difference at the p<0.01 level.

## Discussion

LN remains difficult to manage, and there is a significant need for innovative strategies that sustain long-term remission while enabling a meaningful reduction in GC doses. Observational SLE studies have linked higher omega-3 intake and circulating DHA with diminished disease activity as reflected in SLAQ scores and better patient-reported outcomes, including reduced pain and improved sleep [83–85]; however, randomized controlled trials (RCTs) have yielded mixed results.

Akbar et al. [22] reported variable benefits across nine RCTs that differed in duration (10–52 weeks), sample size (12–85 participants), and dosage (0.54–3.60 g EPA; 0.30–2.25 g DHA). Several studies reported improvements in clinical or serologic markers [86–90], while others observed only transient or no benefit [32,91–93]. A recent RCT using krill oil (4 g/day; 722 mg EPA, 384 mg DHA) showed positive effects in patients with high baseline disease activity (SLEDAI-2K ≥ 9) [16]. Overall, although omega-3 RCTs suggest a potential for modest improvement in SLE disease activity, small sample sizes, inconsistent dosing, varied populations, short follow-up periods, limited assessment of long-term tissue FA incorporation, and suboptimal control of confounding factors hinder clear interpretation. Therefore, the full extent of omega-3s’ benefits for individuals with SLE and LN, along with how they interact with standard treatments, remains unclear.

Because omega-3 supplements are not patent-protected, large, highly optimized randomized controlled trials used to evaluate new drugs are impractical, stressing the importance of rigorous preclinical models to define their therapeutic potential in LN. In this context, the SALN model offers a stringent, translationally relevant platform for evaluating remission-maintenance strategies after immunosuppressive induction. Mechanistically, short-term, repeated cSiO_2_ exposure in NZBWF1 mice combines an environmentally relevant trigger with genetic susceptibility. Inhaled particles are initially phagocytosed by alveolar macrophages (AMs). This results in phagolysosome destabilization, inflammasome activation, and AM death via pyroptosis and necrosis, and ultimately the release of cytokines, chemokines, DAMPs, nuclear antigens, and cSiO_2_ into the local environment [78,94] (Figure 12). These events, together with the long pulmonary half-life of cSiO_2_, create a self-sustaining loop in which dying and activated AMs promote ELS formation in the lung, enhance T- and B-cell recruitment and activation, and accelerate the production of pathogenic AAbs.

**Figure 12.**
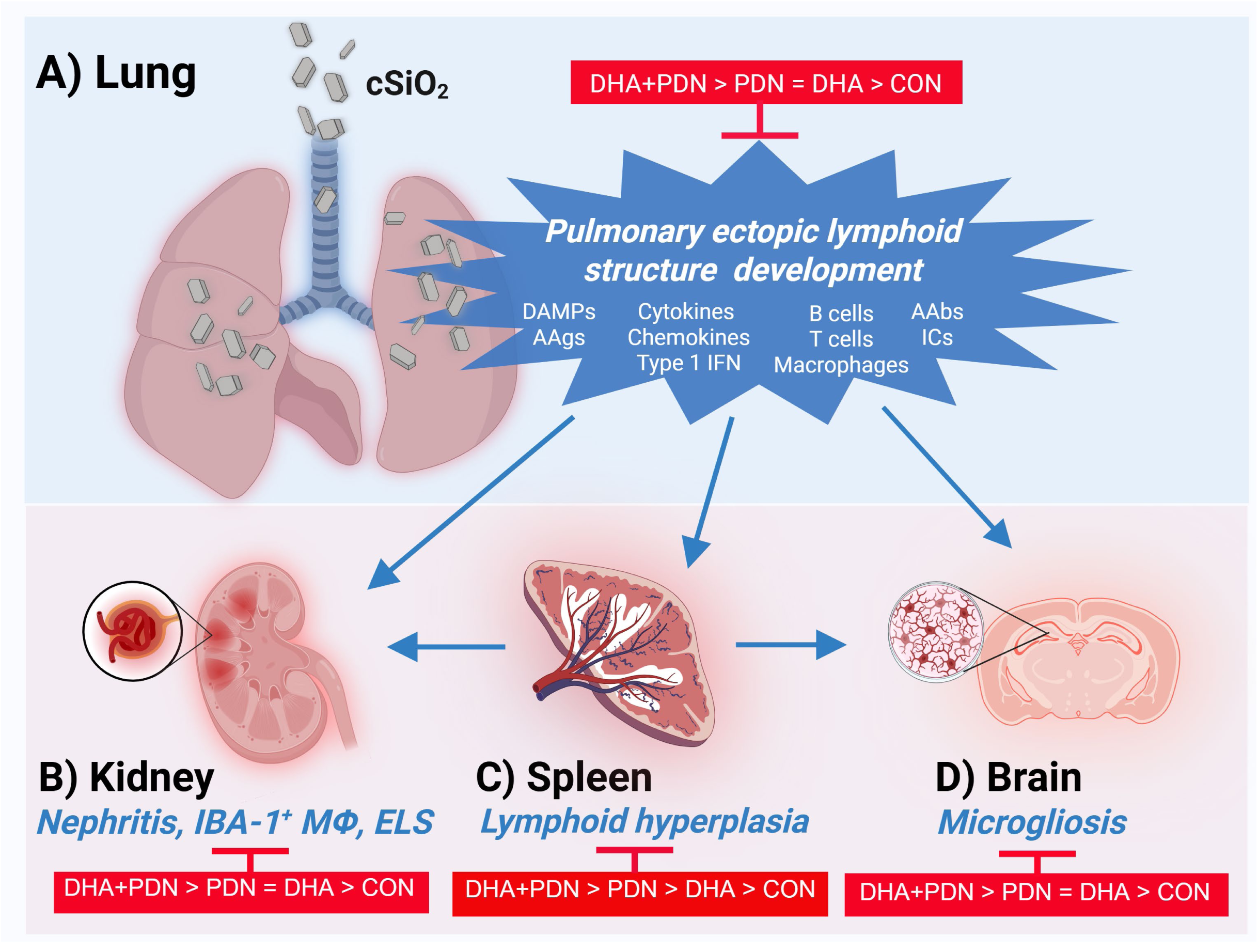
Schematic summarizing how docosahexaenoic acid (DHA) supplementation influences prednisone (PDN) efficacy for the sustained maintenance of cyclophosphamide-induced remission in silica-triggered lupus. Inhaled crystalline silica (cSiO₂) initiates lung cell death and inflammation, inducing damage-associated molecular patterns (DAMPs), autoantigens (AAgs), type I interferons, and chemokines, which together drive robust infiltration by B cells, T cells, and macrophages, leading to ectopic lymphoid structure formation, AAb production, and immune complex (IC) formation. These immunological events in the lung amplify multiorgan pathology, including glomerulonephritis, IBA-1⁺ macrophage accumulation, and lymphoid neogenesis in the kidney, splenic lymphoid hyperplasia, and microgliosis in the brain. Compared to single-agent regimens, combined DHA+PDN therapy exerts superior suppression of multisystem lupus pathology (DHA+PDN > PDN = DHA > control for kidney, lung, and brain; DHA+PDN > PDN > DHA > control for spleen), supporting additive benefits of omega-3 supplementation and long-term maintenance with modest steroid doses to reduce risk of flare and cumulative tissue injury. Figure created using BioRender at https://BioRender.com.

This process drives both pulmonary and systemic autoimmunity and results in severe, synchronized glomerulonephritis (GN) that resembles key features of human SLE and LN [55,56]. Compared with the spontaneous NZBWF1 model, SALN presents a condensed, synchronized multi-organ disease course, with LN developing within 3 months after the final cSiO_2_ exposure. This allows intervention studies to be completed in 7 months, rather than 10 to 12 months in the conventional NZBWF1 model.

Previous investigations employing SALN have shown that either DHA or PDN monotherapy can prevent LN onset [56–58,60–62,95], but these studies mainly focused on disease prevention. The present study is innovative because it endeavors to bridge this translational gap by using SALN to assess DHA as an adjunct therapy to prolong CYC-induced remission, both alone and in combination with a modest, clinically relevant dose of PDN in established LN. We report for the first time that dietary DHA administered post-LN onset during and after immunosuppressive induction prompts significant remodeling of tissue membrane lipid composition. This change is associated with enhanced clinical and histopathological outcomes (Figure 12). Incorporation of DHA and other omega-3 fatty acids into phospholipids, along with a reciprocal depletion of omega-6 species, most notably ARA, indicates robust reprogramming of tissue lipid pools. In mice given DHA+PDN, increases in total omega-3 and decreases in omega-6 were enhanced by 33 percent and 19 percent, respectively, compared with mice fed DHA alone. In the kidney, DHA, particularly when combined with PDN, prolonged CYC-induced remission by delaying proteinuric relapse and maintaining renal structure. This was evidenced by reduced macrophage infiltration, diminished tubular protein deposition, and prevention of ELS formation.

The benefits of DHA combined with PDN were observed beyond the kidney. In the lung, this combination reduced perivascular inflammation, ectopic lymphoid aggregates, and local autoantibody production. Systemically, it decreased splenomegaly and circulating autoantibodies. In the brain, DHA+PDN reduced hippocampal microgliosis and improved neural tissue health, indicating a wider impact on neuroinflammatory pathways in the CNS. Collectively, this evidence indicates that DHA-rich nutrition, particularly when combined with PDN, exerts coordinated immune-modulating effects across multiple organs, enhances the durability of LN remission, and reduces extra-renal inflammation and tissue damage, thereby supporting its ability to augment SLE outcomes more broadly. Thus, adjunctive DHA may enhance conventional two-phase immunosuppressive therapy and is consistent with current rheumatology treatment approaches focused on sustained remission with reduced GC exposure [9].

The organ-level effects observed here in SALN align with the well-known immunomodulatory actions of omega-3 fatty acids. DHA and other omega-3s affect immune responses through interrelated structural, transcriptional, and metabolic pathways [17,18,96,97]. Their incorporation into membrane phospholipids alters lipid raft organization, increases membrane fluidity, and interferes with receptor clustering required for effective signaling through TLRs, TNFR, and IL1R [98]. At the transcriptional level, DHA suppresses NF-κB and activates PPARγ and NRF2, diminishing oxidative stress and decreasing expression of proinflammatory mediators such as IL6 and iNOS [99,100]. Additionally, omega-3s inhibit ARA production from plant-derived LA [101] and competitively inhibit the formation of inflammatory mediators from ARA, such as leukotrienes, prostaglandins, and thromboxanes [96]. DHA also serves as a precursor for proresolving mediators that promote efferocytosis, facilitating the clearance of apoptotic cells and supporting the resolution of chronic inflammation [97]. The evidence presented here is further consistent with preventive studies in NZBWF1 mice exposed to cSiO₂, where DHA elicits broad anti-inflammatory and immunoregulatory actions in the lungs [63]. Following a single intranasal cSiO₂ challenge, control-fed mice showed increased leukocyte influx, cytokine and chemokine levels, dead cell accumulation, and higher AAbs in BALF. In contrast, DHA-enriched diets lessened these responses, including IL1α, IL6, GM-CSF, CCL2, CCL3, and CXCL10, and lowered autoimmune-related gene signatures linked to type I IFN signaling, TLR activation, and lymphocyte activation, with predicted suppression of regulators like TNFα, IL1β, IFNAR, and IFNγ.

The tissue-specific effects of PDN in SALN observed here are in accord with the known immunomodulatory actions of GCs. Treatment with PDN and other synthetic GCs rapidly controls inflammation in SLE and related rheumatic conditions by activating the glucocorticoid receptor (GR), which suppresses NF-κB/AP-1–driven inflammatory pathways in lymphoid and myeloid cells and enhances anti-inflammatory genes like glucocorticoid-induced leucine zipper (GILZ) and dual specificity phosphatase 1 (DUSP1) [102–104]. Non-genomic GR signaling further contributes by inhibiting PLA2 and activating MAPK and cAMP pathways [102,104,105]. Additionally, GCs modify lipid rafts by altering their lipid composition and physical properties, disrupting raft formation in some immune cells while also recruiting GRs and related signaling complexes into flotillin- or caveolin-rich raft regions, thereby affecting kinase signaling and gene regulation [106–108]. Thus, the anti-inflammatory effects of DHA and PDN involve multiple parallel interconnected mechanisms.

Using SALN, we previously found that pretreatment with a moderate PDN dose (14 mg/day HED), taken orally via diet, significantly reduced cSiO₂-induced pulmonary ELS formation, AAb production, and autoimmune gene expression in both the lung and kidney [64]. This treatment also decreased splenomegaly and the severity of glomerulonephritis. Instead of reversing disease signatures, this regimen reduced the magnitude and diversity of transcriptomic changes across organs, showing a general suppression of autoimmune activation. Key adaptive immune modules—including T-cell receptor signaling, antigen presentation, B-cell activation, and Fc-mediated effector pathways—were particularly suppressed, especially in the lung. Overall, these prior findings show that moderate-dose PDN provides widespread but incomplete transcriptional control over the cSiO₂-induced autoimmune process, with innate and stress responses still partly active.

The PDN dose used in the present study was specifically chosen based on its narrow preventive window in SALN: 14 mg/day HED reduces disease progression, 5 mg/day is ineffective, and 46 mg/day causes severe GC-related toxicity without providing a survival benefit [64]. This pattern parallels clinical experience, in which high-dose GCs induce remission but carry significant cumulative toxicity, and highlights that monotherapy suppresses inflammation without restoring immune homeostasis [109]. Based on this prior study, herein we deliberately selected a suboptimal intermediate dose (9.4 mg/day HED), positioned between previously ineffective and minimally effective levels in SALN [64]. Clinically, this dose is near the lower end of moderate-dose GC exposure: in a large SLE cohort, the mean daily PDN-equivalent dose during the first treatment year was 19.4 mg, with about 60% of patients receiving ≥15 mg/day and only 17% maintained below 7.5 mg/day [110]. The 9.4 mg/day HED selected here represents a clinically significant exposure, half of the dose usually employed during the first year of treatment, but still above the current treat-to-target guidelines not to exceed 5 to 7.5 mg/day for SLE and LN maintenance.

Macrophages play a vital role in SLE development, driven by unresolved tissue inflammation, defective removal of cellular debris, and persistent type I IFN signaling [111–113]. In a recent in vitro study using a novel self-renewing AM model derived from embryonic NZBWF1 mouse liver, we determined that DHA synergistically enhances the suppressive effects of low doses of the water-soluble GC dexamethasone (DEX) on LPS-induced type 1 IFN-driven gene expression [27]. When combined, subinhibitory levels of DHA and DEX led to over tenfold greater gene suppression than either agent alone. This combination significantly reprogrammed inflammatory transcriptional networks, decreasing activity in the NFκB, AP1, IRF, and STAT pathways while promoting pathways associated with resolution and M2 polarization. These in vitro mechanistic insights align with the in vivo SALN results discussed here, further supporting DHA as an adjunct therapy to GC and highlighting its potential to enhance treatment outcomes by targeting critical pathogenic pathways in SLE.

Interpretation of these data must consider several limitations. First, the relatively short experimental duration limits conclusions about long-term disease progression and survival, and formal dose–response relationships for both DHA and PDN—individually and together—are yet to be systematically established. Second, the study was intentionally designed to focus on remission maintenance, renal and extrarenal histopathology, and survival rather than a detailed mechanistic analysis of how upstream signaling pathways are affected by DHA and PDN. Third, variations in omega-3 types (DHA vs EPA) and bioavailability (ethyl esters, triglycerides, phospholipids) among formulations on the market may additionally impact treatment effectiveness [16]. Finally, the metabolic and immunological features of SALN may not fully parallel human SLE, so caution is necessary when extrapolating to diverse patient populations. Therefore, to address these limitations, future preclinical research using the SALN model should i) explicitly address dose–response relationships over an extended time period, ii) compare various therapeutic formulations and delivery methods (e.g., structured triglycerides or phospholipid conjugates), and iii) include lipidomic and transcriptomic profiling to understand better how DHA and PDN reprogram immune–metabolic networks across disease. Ultimately, well-designed multicenter clinical trials will be essential to determine the translational relevance and clinical significance of DHA-based adjunctive GC-sparing strategies in SLE and LN.

## Conclusion

This study provides compelling preclinical evidence that DHA supplementation significantly enhances the efficacy of PDN in sustaining CYC-induced remission in LN. By leveraging a robust, translationally relevant model, we demonstrate that DHA exerts immunomodulatory effects that complement glucocorticoid mechanisms, resulting in prolonged renal and extrarenal protection. These findings highlight the potential of omega-3 fatty acids as safe, cost-effective adjuncts to conventional immunosuppressive regimens, offering a strategy to reduce glucocorticoid burden while improving long-term outcomes. If validated in clinical trials, DHA-based combination therapy could redefine LN management by integrating precision nutrition into standard care, thereby advancing treatment paradigms toward sustained remission with minimized toxicity.

## Conflict of Interest

The authors declare that the research was conducted in the absence of any commercial or financial relationships that could be construed as a potential conflict of interest.

## Author Contributions

AA: hypothesis, study design, coordination, animal handling, instillations, urinalysis, necropsy, data curation, data analysis/interpretation, figure preparation, manuscript preparation and submission; OM: animal handling, necropsy, instillations, AAb ELISAs, figure preparation; SJ: animal handling/feeding, injections, urinalysis, diet preparation, sample/data collection, morphometry, figure preparation; JL: animal handling/feeding, diet preparation, data collection; VE: necropsy, BUN assays, figure preparation; JJ: injections, AAb ELISAs; RL: coordinator, instillations, necropsy, sample histology preparation, slide scanning; JW: necropsy, BALF cell counts; JH: necropsy, oversight, lung/kidney/spleen/brain histopathology, semi-quantitative data analysis, manuscript preparation; JP: hypothesis, study design, oversight, funding acquisition, data analysis/ interpretation, manuscript preparation, and submission.

## Funding

This research was funded by NIH ES027353 (AA, JP, JH), DOD HT94252510457 (JP, JH), Lupus Research Alliance Grant 1260473 (AA), and Dr. Robert and Carol Deibel Family Endowment (JP).

## Acknowledgments

The authors thank Amy Porter and Kaila Hogg of the Michigan State University Laboratory for Investigative Histopathology for their assistance with histotechnology.

## Generative Artificial Intelligence (AI)

Generative AI (Perplexity, Perplexity AI, San Francisco, CA, USA) and Grammarly (Grammarly Inc., San Francisco, CA, USA) were used exclusively for language editing and to improve the clarity, readability, and organization of the manuscript. All scientific content, data interpretation, and conclusions were generated by the authors, who carefully reviewed and edited all AI-assisted text and take full responsibility for the integrity and accuracy of the work

## Data Availability Statement

The data supporting the findings of this study are available from the corresponding author upon reasonable request.

## Supplemental Methods

### Immunohistochemistry (IHC)

IHC was conducted as previously described (Pestka et al. 2021, Front. Immunol.12:653464). Briefly, formalin-fixed, paraffin-embedded samples were sectioned, placed on charged slides, and dried overnight at 56°C. The slides were then deparaffinized in xylene and rehydrated using a dilution series of ethanol ending with distilled water. Slides were immersed in tris-buffered saline pH 7.4 (TBS, Scytek Labs, Logan, UT) for 5 minutes to adjust the pH. Following TBS, retrieval protocols were applied according to the requirements of each primary antibody. Endogenous peroxidase activity was blocked for 30 minutes at room temperature in a solution of 3% hydrogen peroxide in methanol (1:4), then rinsed with running tap water, followed by several rinses in distilled water, and a 5-minute incubation in TBS containing Tween 20 (Scytek Labs, Logan, UT). After pretreatment, standard micropolymer complex staining steps were performed at room temperature using the Biocare Medical intelliPATH FLX automated IHC staining instrument. Rinses after each staining step were carried out with TBS autowash buffer containing surfactant (TWB945M, Biocare Medical, Pacheco, CA).

After blocking for nonspecific protein with rodent block M (Biocare Medical, Pacheco, CA) or IntelliPATH background punisher (IP974G20, Biocare Medical, Pacheco, CA) for 20 or 5 minutes, respectively, tissue sections were incubated with primary antibodies (anti-IBA-1, 1:3,000, anti-CD3 1:200, anti-CD45R 1:250) in normal antibody diluent (Scytek Labs, Logan, UT). MicroPolymer Rabbit on Rodent HRP Polymer was used for the CD3 and IBA-1 primary antibodies, and Rat on Mouse HRP Probe and Polymer were used for the CD45R primary antibody. To visualize the target proteins, the substrate Biocare Medical Romulin AEC Chromogen was incubated with the tissue for 5 minutes, then counterstained with CATHE Hematoxylin (Biocare Medical, Pacheco, CA) at a dilution of 1:10 for 1 minute. Afterwards, slides were air-dried, cleared, and mounted with synthetic mounting media.

**Supplementary Table 1.**
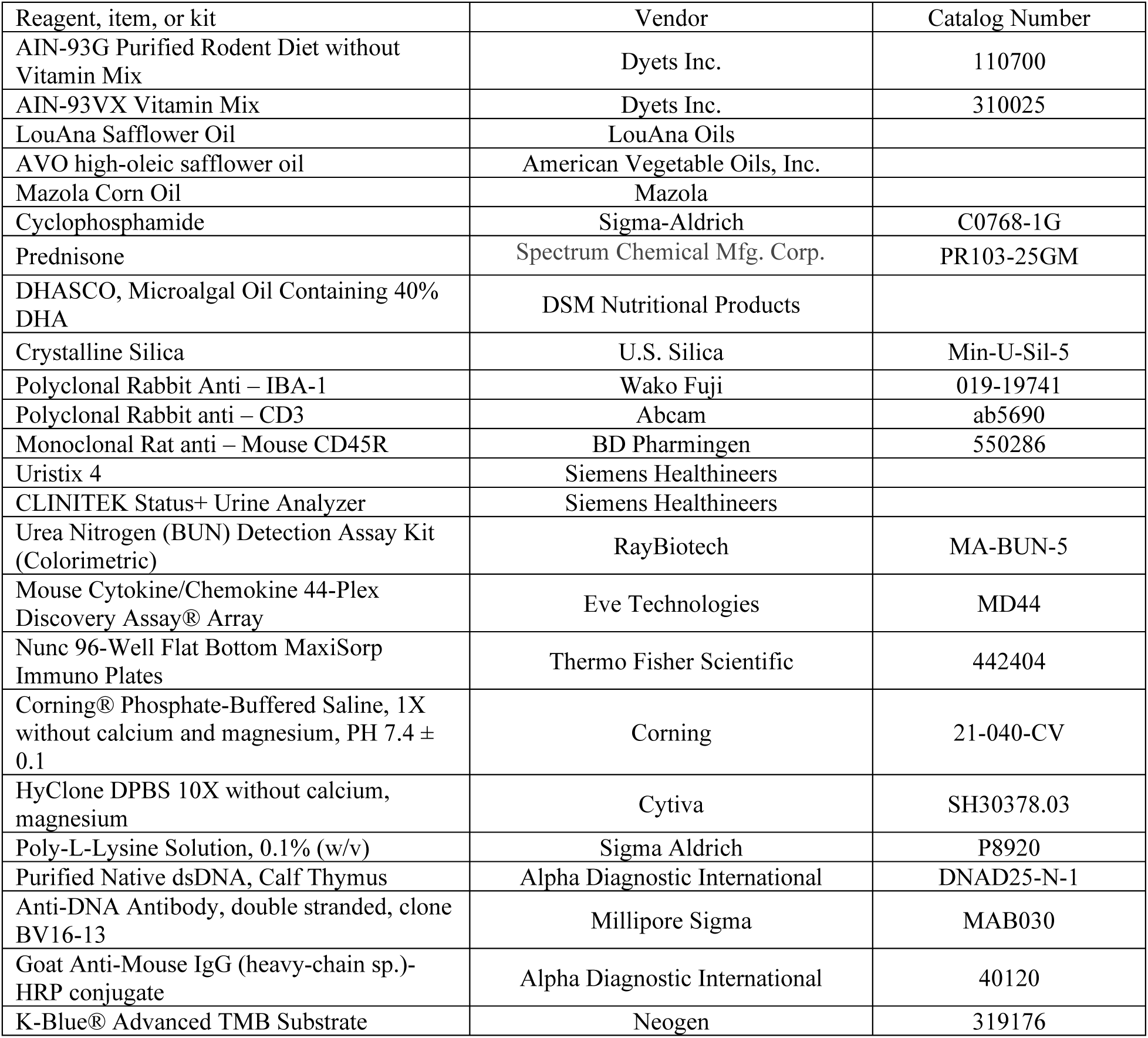
List of key reagents, items, and kits.

**Supplementary Table 2.**
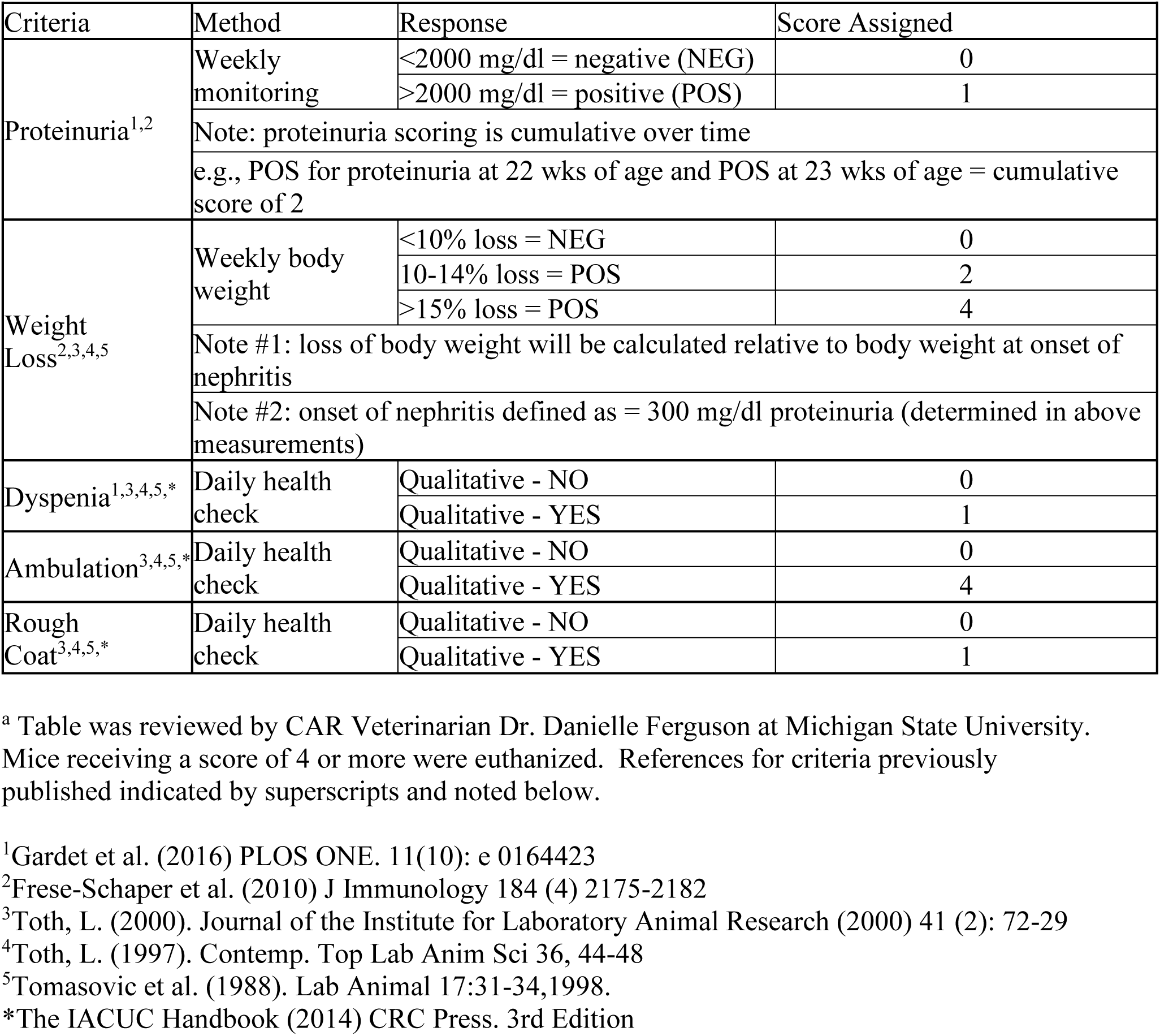
Criteria for defining moribund condition of silica-treated NZBWF1 mice^a^.

**Supplemental Figure 1.**
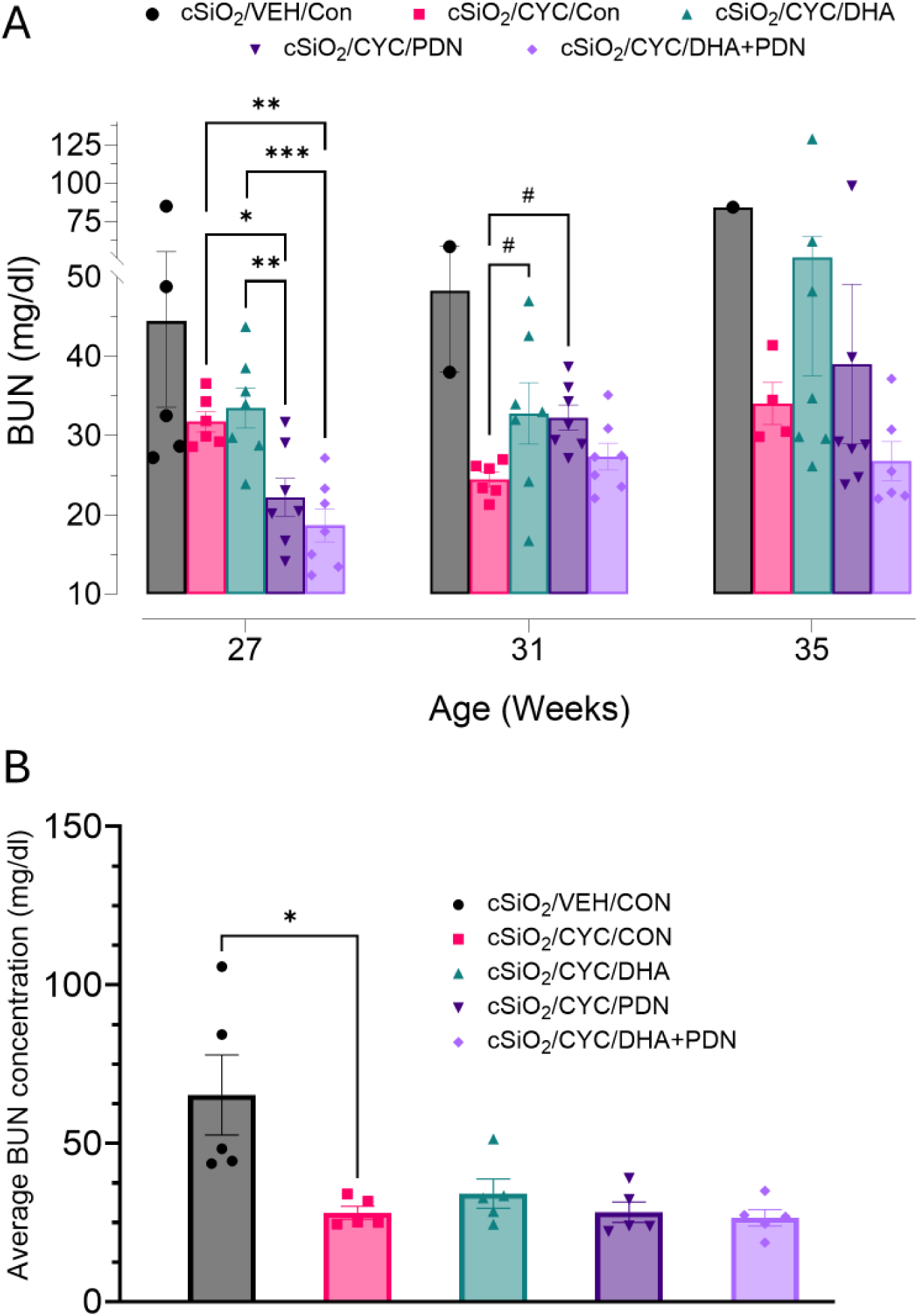
Effects of DHA and PDN monotherapies and co-therapy on CYC-induced reduction of blood urea nitrogen (BUN) in the cSiO_2_-accelerated lupus nephritis mouse model. (A) BUN values at specific time points during the study. (B) Average BUN levels from weeks 27-35 of the study. Symbols: #, *, **, and *** denote significant differences at the p<0.1, p<0.05, p<0.01, and p<0.001 levels, respectively.

## Notes

### Competing Interest Statement

The authors have declared no competing interest.

### Summary of Updates

This version of the manuscript has been revised to correct errors in formatting and punctuation.

## References

[1] Tsokos GC. The immunology of systemic lupus erythematosus. Nat Immunol. 2024;25(8):1332–1343.

[2] Woo JMP, Parks CG, Jacobsen S, Costenbader KH, Bernatsky S. The role of environmental exposures and gene-environment interactions in the etiology of systemic lupus erythematous. J Intern Med. 2022;291(6):755–778.

[3] Caielli S, Wan Z, Pascual V. Systemic lupus erythematosus pathogenesis: Interferon and beyond. Annu Rev Immunol. 2023;41:533–560.

[4] Hoi A, Igel T, Mok CC, Arnaud L. Systemic lupus erythematosus. Lancet. 2024;403(10441):2326–2338.

[5] Anders HJ, Saxena R, Zhao MH, Parodis I, Salmon JE, et al. Lupus nephritis. Nat Rev Dis Primers. 2020;6(1):7.

[6] Desai NB, Gashti C, Whittier WL. "A change is gonna come" to treatment of lupus nephritis: A review. Am J Kidney Dis. 2025.

[7] Fanouriakis A, Kostopoulou M, Andersen J, Aringer M, Arnaud L, et al. Eular recommendations for the management of systemic lupus erythematosus: 2023 update. Ann Rheum Dis. 2024;83(1):15–29.

[8] Mok CC, Teng YKO, Saxena R, Tanaka Y. Treatment of lupus nephritis: Consensus, evidence and perspectives. Nat Rev Rheumatol. 2023;19(4):227–238.

[9] Frangou E, Anders HJ, Bajema IM, Teng YKO, Malvar A, et al. Immunosuppression withdrawal in patients with lupus nephritis: When, how, and for whom will it be safe? J Am Soc Nephrol. 2024;35(7):955–958.

[10] Newman TV, Guo J, Hernandez I. Emerging treatments for lupus nephritis: Health equity considerations in clinical research and coverage. J Manag Care Spec Pharm. 2021;27(10):1500–1502.

[11] Nicastro HL, Vorkoper S, Sterling R, Korn AR, Brown AGM, et al. Opportunities to advance implementation science and nutrition research: A commentary on the strategic plan for nih nutrition research. Transl Behav Med. 2023;13(1):1–6.

[12] Shanmugam VK, Ufret-Vincenty C, Li X, Clayton JA. The inaugural nih-wide strategic plan for autoimmune disease research (fy2026–2030). Arthr Rheumatol.4.

[13] Robinson GA, McDonnell T, Wincup C, Martin-Gutierrez L, Wilton J, et al. Diet and lupus: What do the patients think? Lupus. 2019;28(6):755–763.

[14] Ramessar N, Borad A, Schlesinger N. The effect of omega-3 fatty acid supplementation in systemic lupus erythematosus patients: A systematic review. Lupus. 2022;31(3):287–296.

[15] Salek M, Hosseini Hooshiar S, Salek M, Poorebrahimi M, Jafarnejad S. Omega-3 fatty acids: Current insights into mechanisms of action in systemic lupus erythematosus. Lupus. 2023;32(1):7–22.

[16] Salmon J, Wallace DJ, Rus V, Cox A, Dykas C, et al. Correction of omega-3 fatty acid deficiency and improvement in disease activity in patients with systemic lupus erythematosus treated with krill oil concentrate: A multicentre, randomised, double-blind, placebo-controlled trial. Lupus Sci Med. 2024;11(2).

[17] Calder PC. Omega-3 fatty acids and inflammatory processes: From molecules to man. Biochem Soc Trans. 2017;45(5):1105–1115.

[18] Bodur M, Yilmaz B, Ağagündüz D, Ozogul Y. Immunomodulatory effects of omega-3 fatty acids: Mechanistic insights and health implications. Mol Nutr Food Res. 2025;69(10):e202400752.

[19] Djuricic I, Calder PC. Pros and cons of long-chain omega-3 polyunsaturated fatty acids in cardiovascular health. Annu Rev Pharmacol Toxicol. 2023;63:383–406.

[20] Laguzzi F, Åkesson A, Marklund M, Qian F, Gigante B, et al. Role of polyunsaturated fat in modifying cardiovascular risk associated with family history of cardiovascular disease: Pooled de novo results from 15 observational studies. Circulation. 2024;149(4):305–316.

[21] O’Keefe JH, Tintle NL, Harris WS, O’Keefe EL, Sala-Vila A, et al. Omega-3 blood levels and stroke risk: A pooled and harmonized analysis of 183 291 participants from 29 prospective studies. Stroke. 2024;55(1):50–58.

[22] Akbar U, Yang M, Kurian D, Mohan C. Omega-3 fatty acids in rheumatic diseases: A critical review. J Clin Rheumatol. 2017;23(6):330–339.

[23] Sigaux J, Mathieu S, Nguyen Y, Sanchez P, Letarouilly JG, et al. Impact of type and dose of oral polyunsaturated fatty acid supplementation on disease activity in inflammatory rheumatic diseases: A systematic literature review and meta-analysis. Arthritis Res Ther. 2022;24(1):100.

[24] EFSA Panel on Dietetic Products N, Allergies. Scientific opinion on the tolerable upper intake level of eicosapentaenoic acid (epa), docosahexaenoic acid (dha) and docosapentaenoic acid (dpa). EFSA Journal. 2012;10(7):2815.

[25] Kennedy ET, Luo H, Ausman LM. Cost implications of alternative sources of (n-3) fatty acid consumption in the united states1234. The Journal of Nutrition. 2012;142(3):605S–609S.

[26] Wierenga KA, Strakovsky RS, Benninghoff AD, Rajasinghe LD, Lock AL, et al. Requisite omega-3 hufa biomarker thresholds for preventing murine lupus flaring. Front Immunol. 2020;11:1796.

[27] Heine LK, Nault R, Jackson J, Anderson AN, Harkema JR, et al. Omega-3 fatty acid synergy with glucocorticoid in mouse lupus macrophage model: Targeting pathogenic pathways to reduce steroid dependence. Front Immunol. 2025;16:1646133.

[28] Alexander NJ, Smythe NL, Jokinen MP. The type of dietary fat affects the severity of autoimmune disease in nzb/nzw mice. Am J Pathol. 1987;127(1):106–21.

[29] Bhattacharya A, Lawrence RA, Krishnan A, Zaman K, Sun D, et al. Effect of dietary n-3 and n-6 oils with and without food restriction on activity of antioxidant enzymes and lipid peroxidation in livers of cyclophosphamide treated autoimmune-prone nzb/w female mice. J Am Coll Nutr. 2003;22(5):388–99.

[30] Blok WL, Katan MB, van der Meer JW. Modulation of inflammation and cytokine production by dietary (n-3) fatty acids. J Nutr. 1996;126(6):1515–33.

[31] Chandrasekar B, Troyer DA, Venkatraman JT, Fernandes G. Dietary omega-3 lipids delay the onset and progression of autoimmune lupus nephritis by inhibiting transforming growth factor beta mrna and protein expression. J Autoimmun. 1995;8(3):381–93.

[32] Clark WF, Parbtani A. Omega-3 fatty acid supplementation in clinical and experimental lupus nephritis. Am J Kidney Dis. 1994;23(5):644–7.

[33] Fernandes G, Bysani C, Venkatraman JT, Tomar V, Zhao W. Increased tgf-beta and decreased oncogene expression by omega-3 fatty acids in the spleen delays onset of autoimmune disease in b/w mice. J Immunol. 1994;152(12):5979–87.

[34] Fernandes G, Chandrasekar B, Luan X, Troyer DA. Modulation of antioxidant enzymes and programmed cell death by n-3 fatty acids. Lipids. 1996;31 Suppl(3 SUPPL.):S91–6.

[35] Godfrey DG, Stimson WH, Watson J, Belch JF, Sturrock RD. Effects of dietary supplementation on autoimmunity in the mrl/lpr mouse: A preliminary investigation. Ann Rheum Dis. 1986;45(12):1019–24.

[36] Halade GV, Rahman MM, Bhattacharya A, Barnes JL, Chandrasekar B, et al. Docosahexaenoic acid-enriched fish oil attenuates kidney disease and prolongs median and maximal life span of autoimmune lupus-prone mice. J Immunol. 2010;184(9):5280–6.

[37] Halade GV, Williams PJ, Veigas JM, Barnes JL, Fernandes G. Concentrated fish oil (lovaza(r)) extends lifespan and attenuates kidney disease in lupus-prone short-lived (nzb x nzw)f1 mice. Exp Biol Med 2013;238(6):610–622.

[38] Jolly CA, Fernandes G. Diet modulates th-1 and th-2 cytokine production in the peripheral blood of lupus-prone mice. J Clin Immunol. 1999;19(3):172–8.

[39] Jolly CA, Muthukumar A, Reddy Avula CP, Fernandes G. Maintenance of nf-kappab activation in t-lymphocytes and a naive t-cell population in autoimmune-prone (nzb/nzw)f(1) mice by feeding a food-restricted diet enriched with n-3 fatty acids. Cell Immunol. 2001;213(2):122–33.

[40] Kelley VE, Ferretti A, Izui S, Strom TB. A fish oil diet rich in eicosapentaenoic acid reduces cyclooxygenase metabolites, and suppresses lupus in mrl-lpr mice. J Immunol. 1985;134(3):1914–9.

[41] Kobayashi A, Ito A, Shirakawa I, Tamura A, Tomono S, et al. Dietary supplementation with eicosapentaenoic acid inhibits plasma cell differentiation and attenuates lupus autoimmunity. Front Immunol. 2021;12:650856.

[42] Lim BO, Jolly CA, Zaman K, Fernandes G. Dietary (n-6) and (n-3) fatty acids and energy restriction modulate mesenteric lymph node lymphocyte function in autoimmune-prone (nzb x nzw)f1 mice. J Nutr. 2000;130(7):1657–64.

[43] Cheng Y, Liu L, Ye Y, He Y, Hu W, et al. Roles of macrophages in lupus nephritis. Front Pharmacol. 2024;15:1477708.

[44] Pestka JJ, Vines LL, Bates MA, He K, Langohr I. Comparative effects of n-3, n-6 and n-9 unsaturated fatty acid-rich diet consumption on lupus nephritis, autoantibody production and cd4+ t cell-related gene responses in the autoimmune nzbwf1 mouse. PLoS One. 2014;9(6):e100255.

[45] Prickett JD, Robinson DR, Steinberg AD. A diet enriched with eicosapentaenoic acid suppresses autoimmune nephritis in female (nzb x nzw) f1 mice. Trans Assoc Am Physicians. 1982;95:145–54.

[46] Reddy Avula CP, Lawrence RA, Zaman K, Fernandes G. Inhibition of intracellular peroxides and apoptosis of lymphocytes in lupus-prone b/w mice by dietary n-6 and n-3 lipids with calorie restriction. J Clin Immunol. 2002;22(4):206–19.

[47] Robinson DR, Hirai A, Steinberg AD, Colvin RB. Alleviation of murine autoimmune disease by dietary marine lipids. Adv Prostaglandin Thromboxane Leukot Res. 1987;17B:850–3.

[48] Robinson DR, Prickett JD, Polisson R, Steinberg AD, Levine L. The protective effect of dietary fish oil on murine lupus. Prostaglandins. 1985;30(1):51–75.

[49] Robinson DR, Xu LL, Tateno S, Guo M, Colvin RB. Suppression of autoimmune disease by dietary n-3 fatty acids. J Lipid Res. 1993;34(8):1435–44.

[50] Spurney RF, Ruiz P, Albrightson CR, Pisetsky DS, Coffman TM. Fish oil feeding modulates leukotriene production in murine lupus nephritis. Prostaglandins. 1994;48(5):331–48.

[51] Venkatraman JT, Chu WC. Effects of dietary omega-3 and omega-6 lipids and vitamin e on serum cytokines, lipid mediators and anti-DNA antibodies in a mouse model for rheumatoid arthritis. J Am Coll Nutr. 1999;18(6):602–13.

[52] Watson J, Godfrey D, Stimson WH, Belch JJ, Sturrock RD. The therapeutic effects of dietary fatty acid supplementation in the autoimmune disease of the mrl-mp-lpr/lpr mouse. Int J Immunopharmacol. 1988;10(4):467–71.

[53] Westberg G, Tarkowski A, Svalander C. Effect of eicosapentaenoic acid rich menhaden oil and maxepa on the autoimmune disease of mrl/l mice. Int Arch Allergy Appl Immunol. 1989;88(4):454–61.

[54] Bates MA, Brandenberger C, Langohr I, Kumagai K, Harkema JR, et al. Silica triggers inflammation and ectopic lymphoid neogenesis in the lungs in parallel with accelerated onset of systemic autoimmunity and glomerulonephritis in the lupus-prone nzbwf1 mouse. PLoS One. 2015;10(5):e0125481.

[55] Bates MA, Benninghoff AD, Gilley KN, Holian A, Harkema JR, et al. Mapping of dynamic transcriptome changes associated with silica-triggered autoimmune pathogenesis in the lupus-prone nzbwf1 mouse. Front Immunol. 2019;10:632.

[56] Bates MA, Brandenberger C, Langohr, II, Kumagai K, Lock AL, et al. Silica-triggered autoimmunity in lupus-prone mice blocked by docosahexaenoic acid consumption. PLoS One. 2016;11(8):e0160622.

[57] Bates MA, Akabari P, Gilley KN, Wagner JG, Li N, et al. Dietary docosahexaenoic acid prevents silica-induced development of pulmonary ectopic germinal centers and glomerulonephritis in the lupus-prone nzbwf1 mouse. Front Immunol. 2018;9:2002.

[58] Benninghoff AD, Bates MA, Chauhan PS, Wierenga KA, Gilley KN, et al. Docosahexaenoic acid consumption impedes early interferon- and chemokine-related gene expression while suppressing silica-triggered flaring of murine lupus. Front Immunol. 2019;10:2851.

[59] Gilley KN, Wierenga KA, Chauhuan PS, Wagner JG, Lewandowski RP, et al. Influence of total western diet on docosahexaenoic acid suppression of silica-triggered lupus flaring in nzbwf1 mice. PLoS One. 2020;15(5):e0233183.

[60] Rajasinghe LD, Li QZ, Zhu C, Yan M, Chauhan PS, et al. Omega-3 fatty acid intake suppresses induction of diverse autoantibody repertoire by crystalline silica in lupus-prone mice. Autoimmunity. 2020;53(7):415–433.

[61] Pestka JJ, Akbari P, Wierenga KA, Bates MA, Gilley KN, et al. Omega-3 polyunsaturated fatty acid intervention against established autoimmunity in a murine model of toxicant-triggered lupus. Front Immunol. 2021;12:653464.

[62] Rajasinghe LD, Bates MA, Benninghoff AD, Wierenga KA, Harkema JR, et al. Silica induction of diverse inflammatory proteome in lungs of lupus-prone mice quelled by dietary docosahexaenoic acid supplementation. Front Immunol. 2021;12:781446.

[63] Chauhan PS, Benninghoff AD, Favor OK, Wagner JG, Lewandowski RP, et al. Dietary docosahexaenoic acid supplementation inhibits acute pulmonary transcriptional and autoantibody responses to a single crystalline silica exposure in lupus-prone mice. Front Immunol. 2024;15:1275265.

[64] Heine LK, Benninghoff AD, Ross EA, Rajasinghe LD, Wagner JG, et al. Comparative effects of human-equivalent low, moderate, and high dose oral prednisone intake on autoimmunity and glucocorticoid-related toxicity in a murine model of environmental-triggered lupus. Front Immunol. 2022;13:972108.

[65] Reeves PG, Nielsen FH, Fahey GC, Jr. Ain-93 purified diets for laboratory rodents: Final report of the american institute of nutrition ad hoc writing committee on the reformulation of the ain-76a rodent diet. J Nutr. 1993;123(11):1939–51.

[66] Buttgereit F, Da Silva JAP, Boers M, Burmester GR, Cutolo M, et al. Standardised nomenclature for glucocorticoid dosages and glucocorticoid treatment regimens: Current questions and tentative answers in rheumatology. Annals of the Rheumatic Diseases. 2002;61(8):718–722.

[67] OSHA. Osha’s final rule to protect workers from exposure to respirable crystalline silica2016 [cited 16286 - 16890 p.]. https://www.federalregister.gov/documents/2016/03/25/2016-04800/occupational-exposure-to-respirable-crystalline-silica

[68] Bankhead P, Loughrey MB, Fernández JA, Dombrowski Y, McArt DG, et al. Qupath: Open source software for digital pathology image analysis. Sci Rep. 2017;7(1):16878.

[69] Meyerholz DK, Beck AP. Fundamental concepts for semiquantitative tissue scoring in translational research. ILAR J. 2018;59(1):13–17.

[70] McDonald OF, Wagner JG, Lewandowski RP, Heine LK, Estrada V, et al. Impact of soluble epoxide hydrolase inhibition on silica-induced pulmonary fibrosis, ectopic lymphoid neogenesis, and autoantibody production in lupus-prone mice. Inhal Toxicol. 2024;36(7-8):442–460.

[71] Harris WS. Recent studies confirm the utility of the omega-3 index. Curr Opin Clin Nutr Metab Care. 2025;28(2):91–95.

[72] Ohsawa K, Imai Y, Sasaki Y, Kohsaka S. Microglia/macrophage-specific protein iba1 binds to fimbrin and enhances its actin-bundling activity. J Neurochem. 2004;88(4):844–56.

[73] Wang M, Rajkumar S, Lai Y, Liu X, He J, et al. Tertiary lymphoid structures as local perpetuators of organ-specific immune injury: Implication for lupus nephritis. Front Immunol. 2023;14:1204777.

[74] Masum MA, Ichii O, Elewa YHA, Otani Y, Namba T, et al. Vasculature-associated lymphoid tissue: A unique tertiary lymphoid tissue correlates with renal lesions in lupus nephritis mouse model. Front Immunol. 2020;11:595672.

[75] Dorraji SE, Kanapathippillai P, Hovd AK, Stenersrød MR, Horvei KD, et al. Kidney tertiary lymphoid structures in lupus nephritis develop into large interconnected networks and resemble lymph nodes in gene signature. Am J Pathol. 2020;190(11):2203–2225.

[76] Harbeck RJ, Launder T, Staszak C. Mononuclear cell pulmonary vasculitis in nzb/w mice. Ii. Immunohistochemical characterization of the infiltrating cells. Am J Pathol. 1986;123(2):204–11.

[77] Staszak C, Harbeck RJ. Mononuclear-cell pulmonary vasculitis in nzb/w mice. I. Histopathologic evaluation of spontaneously occurring pulmonary infiltrates. Am J Pathol. 1985;120(1):99–105.

[78] Wierenga KA, Harkema JR, Pestka JJ. Lupus, silica, and dietary omega-3 fatty acid interventions. Toxicol Pathol. 2019;47(8):1004–1011.

[79] Ye YL, Chiang BL. Reconstitution of severe combined immunodeficient mice with spleen cells from autoimmune nzbxnzw f1 mice. Clin Exp Rheumatol. 1998;16(1):33–7.

[80] Nikolopoulos D, Manolakou T, Polissidis A, Filia A, Bertsias G, et al. Microglia activation in the presence of intact blood-brain barrier and disruption of hippocampal neurogenesis via il-6 and il-18 mediate early diffuse neuropsychiatric lupus. Ann Rheum Dis. 2023;82(5):646–657.

[81] Nomura A, Noto D, Murayama G, Chiba A, Miyake S. Unique primed status of microglia under the systemic autoimmune condition of lupus-prone mice. Arthritis Res Ther. 2019;21(1):303.

[82] Rodríguez-Gómez JA, Kavanagh E, Engskog-Vlachos P, Engskog MKR, Herrera AJ, et al. Microglia: Agents of the cns pro-inflammatory response. Cells. 2020;9(7):1717.

[83] Charoenwoodhipong P, Harlow SD, Marder W, Hassett AL, McCune WJ, et al. Dietary omega polyunsaturated fatty acid intake and patient-reported outcomes in systemic lupus erythematosus: The michigan lupus epidemiology and surveillance program. Arthritis Care Res (Hoboken). 2020;72(7):874–881.

[84] Vordenbaumen S, Sokolowski A, Kutzner L, Rund KM, Dusing C, et al. Erythrocyte membrane polyunsaturated fatty acid profiles are associated with systemic inflammation and fish consumption in systemic lupus erythematosus: A cross-sectional study. Lupus. 2020;29(6):554–559.

[85] Gilley KF, JI; Zick, SM; Li, K;, Wang LM, W;, Mccune WJ, R; Herndon-Fenton, S; Hassett, A; Barbour, K; Pestka, J.J.; Somers, EC;. Serum fatty acid profiles in systemic lupus erythematosus and patient reported outcomes: The michigan lupus epidemiology & surveillance (miles) program. Frontiers in Immunology. 2024;15.

[86] Arriens C, Hynan LS, Lerman RH, Karp DR, Mohan C. Placebo-controlled randomized clinical trial of fish oil’s impact on fatigue, quality of life, and disease activity in systemic lupus erythematosus. Nutr J. 2015;14:82.

[87] Duffy EM, Meenagh GK, McMillan SA, Strain JJ, Hannigan BM, et al. The clinical effect of dietary supplementation with omega-3 fish oils and/or copper in systemic lupus erythematosus. J Rheumatol. 2004;31(8):1551–6.

[88] Lozovoy MA, Simão AN, Morimoto HK, Scavuzzi BM, Iriyoda TV, et al. Fish oil n-3 fatty acids increase adiponectin and decrease leptin levels in patients with systemic lupus erythematosus. Mar Drugs. 2015;13(2):1071–83.

[89] Walton AJ, Snaith ML, Locniskar M, Cumberland AG, Morrow WJ, et al. Dietary fish oil and the severity of symptoms in patients with systemic lupus erythematosus. Ann Rheum Dis. 1991;50(7):463–6.

[90] Wright SA, O’Prey FM, McHenry MT, Leahey WJ, Devine AB, et al. A randomised interventional trial of omega-3-polyunsaturated fatty acids on endothelial function and disease activity in systemic lupus erythematosus. Ann Rheum Dis. 2008;67(6):841–8.

[91] Westberg G, Tarkowski A. Effect of maxepa in patients with sle. A double-blind, crossover study. Scand J Rheumatol. 1990;19(2):137–43.

[92] Bello KJ, Fang H, Fazeli P, Bolad W, Corretti M, et al. Omega-3 in sle: A double-blind, placebo-controlled randomized clinical trial of endothelial dysfunction and disease activity in systemic lupus erythematosus. Rheumatol Int. 2013;33(11):2789–96.

[93] Clark WF, Parbtani A, Huff MW, Reid B, Holub BJ, et al. Omega-3 fatty acid dietary supplementation in systemic lupus erythematosus. Kidney Int. 1989;36(4):653–60.

[94] Favor OK, Pestka JJ, Bates MA, Lee KSS. Centrality of myeloid-lineage phagocytes in particle-triggered inflammation and autoimmunity. Front Toxicol. 2021;3:777768.

[95] Heine LK, Scarlett T, Wagner JG, Lewandowski RP, Benninghoff AD, et al. Crystalline silica-induced pulmonary inflammation and autoimmunity in mature adult nzbw/f1 mice: Age-related sensitivity and impact of omega-3 fatty acid intervention. Inhal Toxicol. 2024;36(2):106–123.

[96] Calder PC. Omega-3 polyunsaturated fatty acids and inflammatory processes: Nutrition or pharmacology? Br J Clin Pharmacol. 2013;75(3):645–62.

[97] Djuricic I, Calder PC. Beneficial outcomes of omega-6 and omega-3 polyunsaturated fatty acids on human health: An update for 2021. Nutrients. 2021;13(7):2421.

[98] Pennington ER, Virk R, Bridges MD, Bathon BE, Beatty N, et al. Docosahexaenoic acid controls pulmonary macrophage lipid raft size and inflammation. J Nutr. 2024;154(6):1945–1958.

[99] Wierenga KA, Riemers FM, Westendorp B, Harkema JR, Pestka JJ. Single cell analysis of docosahexaenoic acid suppression of sequential lps-induced proinflammatory and interferon-regulated gene expression in the macrophage. Front Immunol. 2022;13:993614.

[100] Chang HY, Lee HN, Kim W, Surh YJ. Docosahexaenoic acid induces m2 macrophage polarization through peroxisome proliferator-activated receptor gamma activation. Life Sci. 2015;120:39–47.

[101] Ralston JC, Matravadia S, Gaudio N, Holloway GP, Mutch DM. Polyunsaturated fatty acid regulation of adipocyte fads1 and fads2 expression and function. Obesity. 2015;23(4):725–728.

[102] Furst R, Schroeder T, Eilken HM, Bubik MF, Kiemer AK, et al. Mapk phosphatase-1 represents a novel anti-inflammatory target of glucocorticoids in the human endothelium. FASEB J. 2007;21(1):74–80.

[103] Cain DW, Cidlowski JA. Immune regulation by glucocorticoids. Nat Rev Immunol. 2017;17(4):233–247.

[104] Vandewalle J, Luypaert A, De Bosscher K, Libert C. Therapeutic mechanisms of glucocorticoids. Trends Endocrinol Metab. 2018;29(1):42–54.

105. [105]Cain DW, Cidlowski JA. Immune regulation by glucocorticoids. Nature Publishing Group; 2017. p. 233–247.

[106] Atkovska K, Klingler J, Oberwinkler J, Keller S, Hub JS. Rationalizing steroid interactions with lipid membranes: Conformations, partitioning, and kinetics. ACS Cent Sci. 2018;4(9):1155–1165.

[107] Yamagata S, Tomita K, Sano H, Itoh Y, Fukai Y, et al. Non-genomic inhibitory effect of glucocorticoids on activated peripheral blood basophils through suppression of lipid raft formation. Clin Exp Immunol. 2012;170(1):86–93.

[108] Van Laethem F, Liang X, Andris F, Urbain J, Vandenbranden M, et al. Glucocorticoids alter the lipid and protein composition of membrane rafts of a murine t cell hybridoma. J Immunol. 2003;170(6):2932–9.

[109] Mok TC, Mok CC. Glucocorticoid in systemic lupus erythematosus: The art beyond science. Expert Rev Clin Immunol. 2025;21(5):543–553.

[110] Birt JA, Wu J, Griffing K, Bello N, Princic N, et al. Corticosteroid dosing and opioid use are high in patients with sle and remain elevated after belimumab initiation: A retrospective claims database analysis. Lupus Sci Med. 2020;7(1).

[111] Li Y, Lee PY, Reeves WH. Monocyte and macrophage abnormalities in systemic lupus erythematosus. Arch Immunol Ther Exp (Warsz). 2010;58(5):355–64.

[112] Udalova IA, Mantovani A, Feldmann M. Macrophage heterogeneity in the context of rheumatoid arthritis. Nat Rev Rheumatol. 2016;12(8):472–85.

[113] Northcott M, Jones S, Koelmeyer R, Bonin J, Vincent F, et al. Type 1 interferon status in systemic lupus erythematosus: A longitudinal analysis. Lupus Sci Med. 2022;9(1):e000625.

[114] USFDA. Gras notice (grn) no. 553: Algal oil (40% docosahexaenoic acid) derived from schizochytrium sp. Silver Spring, MD: U.S. Food and Drug Administration, Center for Food Safety and Applied Nutrition; 2015.

[115] Lehman AJ. Approximate relation of parts per million in the diet to mg/kg bw/day. Association of Food and Drug Officials Quarterly Bulletin. 1954 18.

[116] FDA. Estimating the safe starting dose in clinical trials for therapeutics in adult healthy volunteers.: U.S. Department of Health and Human Services 2005. Available from: https://www.fda.gov/regulatory-information/search-fda-guidance-documents/estimating-maximum-safe-starting-dose-initial-clinical-trials-therapeutics-adult-healthy-volunteers

[117] Fryar C, Carroll M, Gu Q, Afful J, Ogden C. Anthropometric reference data for children and adults: United states, 2015–2018. National Center for Health Statistics Vital Health Stat 2021;3(46).

